# PARP inhibitors affect the replisome and replication fork progression during a single S phase

**DOI:** 10.1101/2025.02.20.639283

**Authors:** Lina-Marie Briu, Chrystelle Maric, Nicolas Valentin, Nicolas Panara, Guillaume Chevreux, Giuseppe Baldacci, Jean-Charles Cadoret

## Abstract

Poly(ADP-ribosyl)ation (PARylation) is a protein modification mostly synthesised and degraded by PARP1/2 and PARG enzymes, respectively. PARylation is involved in many covalent and non-covalent protein-protein interactions in the nucleus, making it a powerful form of molecular signaling in DNA metabolism. PARP inhibitors (PARPi) have shown efficacy in the treatment of Homologous Recombination (HR)-deficient cancers, yet the full range of molecular mechanisms underlying the activity of these drugs is not fully understood. Here, we decipher the direct consequences of PARPi-induced loss of PARylation on DNA replication. First, PARPi treatment during a single S phase induces replicative stress, delays S phase progression and causes genome-wide replication timing changes of disease-associated regions. These DNA replication alterations appear to be caused by an accumulation of SSBs in the DNA replicative template when PARylation is inhibited, which then seems to lead to one-ended DSBs and fork collapse during S phase. Second, PARPi deplete FANCD2-I, BRCA, monoubiquitinated PCNA and RAD18 proteins from the replisomes. The two PARPi tested then appear to modulate the choice of fork restart mechanisms and influence replisome dynamics in different ways. Taken together, our study highlights the common and unique primary effects of two PARPi on unperturbed DNA replication in human cells.

**GRAPHICAL ABSTRACT:** 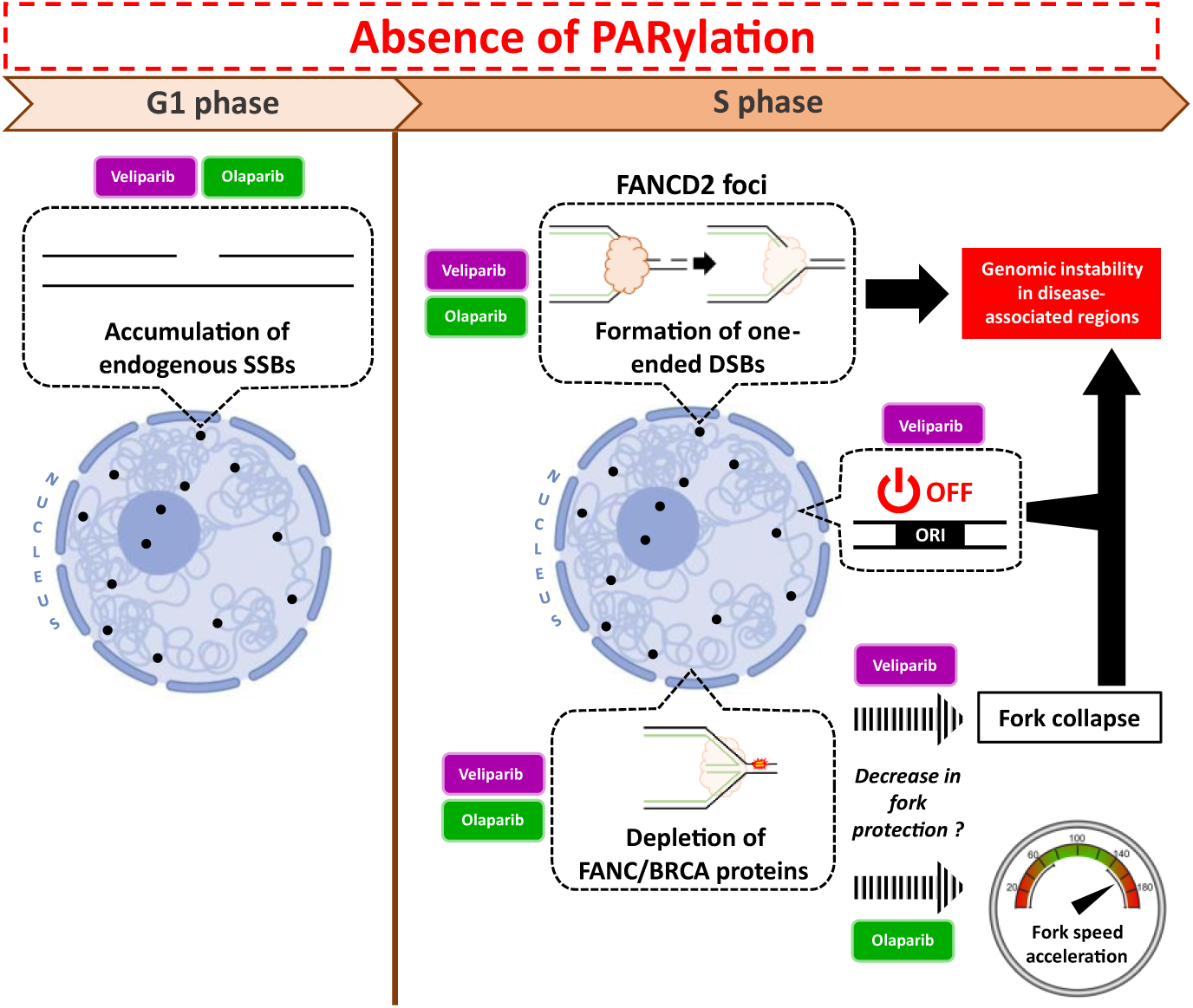

## INTRODUCTION

PolyADP-ribosylation (PARylation) is a post-translational modification of proteins, characterized by the addition of ADP-ribose moieties onto proteins. Nuclear PARylation is primarily synthesized by poly(ADP-ribose) polymerases (PARPs) 1 and 2 from NAD+, and subsequently degraded by poly(ADP-ribose) glycohydrolase (PARG) enzymes. The simultaneous actions of PARP1/2 and PARG result in PARylation being a highly dynamic process, which makes it an effective form of molecular signaling, particularly in the nucleus of human cells. PARylation can form long polymers covalently attached to proteins, which create a highly negatively-charged environment. This environment modulates the proper enzymatic activity of the modified proteins or enables stable interactions between PARylated proteins and PAR-binding proteins. The recruitment of proteins via autoPARylated-PARP1 has been extensively described in the context of DNA damage detection, DNA repair and also in chromatin compaction and transcription regulation [for reviews, 1–3].

It is now well-established that genetic instability in tumor cells occurs mainly during S phase as a consequence of altered DNA replication [for a review, 4]. PARP1 and PARylation are frequently assumed to play a role in the DNA replication process, particularly in the presence of replicative stress. Single-stranded DNA and stalled forks are recognized by PARP1, which then activates its enzymatic activity [5,6]. PARP1 activity has been demonstrated to promote ATR signaling in several studies [7–9]. PARP1 activity promotes fork reversal and stabilizes replication forks by inhibiting the activity of RECQ1, thereby preventing premature fork restart [10,11]. In addition, PARP1 activity has been shown to protect stalled forks from excessive MRE11-dependent nucleolytic degradation [12] as well as to promote “beneficial” MRE11-dependent resection of DNA to mediate recombination repair and restart of collapsed forks [5]. It has recently been demonstrated that PARP1 activity is also required in an alternative process of Okazaki fragment resolution when the canonical process mediated by FEN1 is deficient [13]. However, most of these studies were conducted in a specific mutation context or in cells that had been pre-treated with genotoxic agents. Consequently, they do not provide insight into the role of PARylation in non-perturbed DNA replication. Interestingly, some DNA replication proteins are known to be PARylated [14–20], others to interact with PARP1/2 [21–26] or to bind PARylation [26,27]. Moreover, double-knockout of PARP1 and PARP2 or knockout of PARG in early mouse embryos leads to lethality [28,29], and PARP1 is more highly expressed in proliferative and immature cells across 21 different monkey tissues [30]. This highlights the importance of these enzymes in highly proliferative cells and suggests a pivotal role for PARP1/2 and PARG enzymes in maintaining a balanced PARylation level during DNA replication.

Twenty years ago, it was discovered that the inhibition of PARP1/2 specifically induces cell death of BRCA1 and BRCA2 deficient cells by increasing genomic instability [31,32]. This discovery prompted the development of PARP inhibitors (PARPi) for clinical use, and so far, some of these inhibitors are currently employed in the treatment of breast and ovarian cancers with BRCA1-2 mutations. Although it is now well established that the toxicity of PARP inhibitors requires DNA replication [33, for a review, 34], the major mechanism underlying this toxicity remains a topic of debate [for a review on the different models of PARP inhibitors toxicity, 35] and resistance to these drugs frequently appears in the clinic [for a review, 36]. Recent studies in human cell lines have demonstrated that PARP inhibition alone affects DNA replication and induces genomic instability in human cells. PARP inhibition for either a brief or extended period accelerates fork replication speed, coupled with induction of replication stress and activation of the DNA damage response [37]. This observation has since been confirmed in other work [at least in 33,38]. Among the PARPi, olaparib appears to increase fork speed by activating repriming, as recently published [39], but the exact molecular mechanism remains unclear. Additionally, olaparib has been shown to induce post-replicative single-strand gaps, either by activating the translesion polymerase PRIMPOL [33,40] or by causing unresolved Okazaki fragments [13,41]. These gaps will be toxic in the subsequent S phase, particularly if they become trapped by PARP1 [discussed in 33,37], although no direct evidence has yet been published. Despite the fact that this gap effect has only been demonstrated in the context of olaparib, it seems to explain the sensitivity of HR-deficient cells to PARP inhibitors, since they are devoid of efficient DNA repair pathway and/or fork protection that can effectively address gaps and/or DNA-Protein Crosslinks (DPCs) in the template DNA. This emerging “gap model” seems to provide a compelling rationale for fully reflecting the tumor response to PARP inhibitors in the clinic [42; and for recent reviews 34,43]. However, this “gap model” of PARPi toxicity describes only trans-cell cycle genotoxic effects of PARPi [33], as the post-replicative gaps induced by these inhibitors will challenge DNA replication in the subsequent S phase.

PARP1 and PARP2 enzymes are also involved in single-strand break repair (SSBR) by detecting SSBs and promoting recruitment of SSBR proteins such as XRCC1, PNKP and LIG3 [for a recent review 44]. PARylation is therefore thought to facilitate the base excision repair (BER) process by promoting the repair of the transient SSB generated during the removal of the damaged base. Therefore, the classic model for PARP inhibitor toxicity in HR-deficient cells was that PARP inhibition results in an increase in SSBs that are converted into one-ended DSBs during S phase, which remain unrepaired and cause fork collapse and genomic toxicity in cells lacking HR proteins [31,32]. However, although some studies have shown that PARP inhibitors delay or prevent the repair of induced SSB or BER intermediates [45,46], other studies have shown that inhibition of PARP enzymes or genetic inactivation of *PARP1* alone does not result in the accumulation of SSBs [reviewed in 47]. In addition, experimental models using genetic inactivation of PARylation or prolonged treatment with PARP inhibitors may involve multiple cell cycles, making it impossible to distinguish between the toxic effect on DNA replication arising from PARPi-induced breaks behind the replication forks versus that arising from breaks induced ahead of replication forks. The primary effects of PARylation deficiency on DNA replication are therefore still far from being understood.

In this study, we aim to understand precisely how two PARP inhibitors, olaparib and veliparib, directly influence the dynamics of DNA replication during a single S phase in human cells. We show that these two inhibitors delay the S phase progression, induce replicative stress and decrease DNA synthesis during a single S phase, indicating that the source of these PARPi-associated replication defects occurs during this S phase. We then demonstrate that absence of PARylation leads to the accumulation of SSBs in the replicative template DNA and changes the composition in proteins of the replisome. These two cis effects, when combined, unravel the primary sources of toxicity induced by PARP inhibitors during a single S phase. Finally, we highlight differential effects between olaparib and veliparib on replication fork progression, replication timing program and potential consequences of PARP1-2 inhibition on genome stability. We show that olaparib increases fork speed and has little effect on origin activity and genome replication timing program. In contrast, veliparib has no effect on fork speed but induces fork stalling, decreases origin firing and delays replication timing in many fragile regions enriched in genes associated with synapse organization. These findings offer novel insights into the biology of PARylation during the S phase, and unravel the earliest common and unique sources of toxicity induced by two PARP inhibitors on DNA replication.

## MATERIAL AND METHODS

### Cell cultures

K562 cells were cultured in Roswell Park Memorial Institute (RPMI) 1640 Medium, GlutaMAX™ (Thermo Fisher Scientific, #61870010) supplemented with 10% decomplemented fetal bovine serum (FBS) (Thermo Fisher Scientific, #A5256701), 200U/mL penicillin and 200µg/mL streptomycin (Thermo Fisher Scientific, #15140122). RPE1 cells were cultured in Dulbecco’s modified Eagle’s Medium (DMEM)/F-12, GlutaMAX™ (Thermo Fisher Scientific, #31331028) supplemented with 10% decomplemented FBS, 100U/mL penicillin and 100µg/mL streptomycin. Cells were cultured in a humidified incubator at 37°C with 5% CO2. All experiments were conducted on cells in the exponential growth phase.

### Inhibitors and drugs

Olaparib (Euromedex, #SE-S1060-10MM-1ML) and veliparib (CliniSciences, #A3002) were used as PARP inhibitors. The inhibitors were dissolved in dimethylsulfoxide (DMSO) at concentrations of 10mM (olaparib) or 50mM (veliparib) for storage purposes, with the intention of performing cell treatment at a 1:1000 dilution. Hydroxyurea (HU) (Merck, #H8627) and Hydrogen peroxide (H2O2) (Merck, #88597) were used as positive controls for DNA damage induction. Methyl methanesulfonate (MMS) (Merck, #129925) was used as a positive control for PARP1/2 trapping.

### Synchronization of cells with Elutriation

3×10^8^ asynchronous and exponentially growing K562 cells were pelleted and resuspended in 60mL of dPBS 1X (Thermo Fisher Scientific, #14190094), 1% decomplemented FBS and 1mM EDTA. Cells were loaded in the elutriator rotor (JE-5.0 Beckman Coulter) at a constant speed of 2,500 rpm and a 12mL/min flow rate. Successive increases of flow rate allow the collection of different cell fractions corresponding to early G1 (28mL/min), later G1 (34mL/min), early S (40mL/min) and G2 (82mL/min) phases. 1×10^6^ cells from each fraction were analyzed by flow cytometry to check cell cycle progression.

### Cell Cycle Analysis

K562 cells were fixed with 30% dPBS 1X and 70% cold ethanol and then counterstained with propidium iodide (50μg/ml) (Thermo Fisher Scientific, #P3566) and treated with RNase A (0,5mg/ml) (Merck, #10109169001) for 30 minutes. Samples were acquired using: 1) the Accuri C6 Plus (BD Biosciences) using a 488 nm laser for scatter measurements, including Forward Scatter (FSC) and Side Scatter (SSC), as well as Propidium Iodide (PI) excitation. The PI fluorescence signals were collected through a 585/40 nm band-pass filter. 2) or the FACSAria Fusion (BD Biosciences) with BD FACSDiva Software Version 9.4 (BD Biosciences). The instrument configuration included a 488 nm laser for scatter measurements (FSC and SSC) and a 561 nm laser for PI excitation. The PI fluorescence signals were collected using a 610/20 nm band-pass filter. After the elimination of cell debris and doublets, at least 10,000 cells were analyzed to determine the percentage of cells in each phase of the cell cycle. Flow cytometry data were analyzed using FlowJo V10.10 (BD Biosciences), followed by a quality control and data cleaning step using the PeacoQC algorithm [48] to ensure accuracy and reliability of the results. Cell cycle was analyzed in two independent experiments. The percentage of cells in each phase of the cell cycle was obtained using the Watson Pragmatic algorithm of the Cell Cycle platform in FlowJo V10.10 (BD Biosciences).

### EdU incorporation in nascent DNA

K562 or RPE1 cells were incubated with 10 µM EdU (5-ethynyl-2-deoxyuridine) in culture media for 90 min before the end of the drug treatment. Click-it reactions were performed using the Click-iT™ EdU Alexa Fluor™ 647 Flow Cytometry Assay Kit (Thermo Fisher Scientific, #C10424) or the iClick™ EdU Andy Fluor™ 647 Flow Cytometry Assay Kit (ABP Biosciences, #A008) according to the manufacturer’s recommendations. Cells were then counterstained with propidium iodide (Thermo Fisher Scientific, #P3566) and treated with RNase A (Merck, #10109169001) for 30 minutes. Samples were acquired using: 1) the CyAn ADP LX flow cytometer (Beckman Coulter) with Summit V4.3.04 software (Beckman Coulter). A 488 nm laser was utilized for scatter measurements (FSC and SSC) and (PI) excitation. Additionally, a 635 nm laser was employed for AlexaFluor 647 (AF647) excitation. The fluorescence signals for PI and AF647 were collected using a 613/20 nm band-pass filter and a 665/20 nm band-pass filter, respectively. 2) or the FACSAria Fusion (BD Biosciences) equipped with BD FACSDiva Software Version 9.4 (BD Biosciences). The setup included a 488 nm laser for scatter measurements (FSC and SSC), a 561 nm laser for PI excitation, and a 633 nm laser for AF647 excitation. The fluorescence signals for PI and AF647 were collected using a 610/20 nm band-pass filter and a 670/45 nm band-pass filter, respectively. After the elimination of cell debris and doublets, at least 10,000 cells were analyzed to determine the percentage of cells in each phase of the cell cycle. Flow cytometry data were analyzed using FlowJo V10.10 (BD Biosciences), followed by a quality control and data cleaning step using the PeacoQC algorithm [48] to ensure accuracy and reliability of the results. The EdU incorporation in S phase cells was analyzed in three independent experiments.

### Protein total extraction

About 5×10^6^ K562 cells were used for each condition to extract total proteins. Harvested cells were washed twice with cold dPBS 1X. Cell pellets were resuspended in protein lysis buffer (25 mM Tris-HCl pH 7.5, 100 mM NaCl, 1 mM EDTA, 1 mM EGTA, 0.5% NP40 (Merck, #492016), 1% Triton X-100 (Merck, #T8787) supplemented with 1X cOmplete™ Protease Inhibitor Cocktail (Merck, #11836145001) and Phosphatase Inhibitor Cocktail Set II (Merck, #524625), and incubated for 30 minutes in ice with regular vortexing. Samples were then centrifuged for 10 minutes at 11,000 rpm (HITACHI KOKI Himac CT15E Centrifuge). Supernatants were collected as the total protein fraction. The protein concentrations were measured with a Bradford protein assay. About 20µg proteins were denatured in 2X Laemmli buffer (Merck, #S3401) at 95°C for 5 minutes. In **Figures 2B** and **S4G**, three independent experiments were conducted.

### Chromatin fractionation

3×10^6^ K562 cells were used for each condition to extract chromatin bound proteins. Harvested cells were washed twice with cold dPBS 1X. Cell pellets were lysed in Buffer A (10mM Hepes pH 7.9, 10 mM KCl, 1.5 mM MgCl2, 0.34 M sucrose, 10% glycerol, 1 mM DTT, 10 mM NaF, 1X cOmplete™ Protease Inhibitor Cocktail (Merck, #11836145001), 1X Phosphatase Inhibitor Cocktail Set II (Merck, #524625) and 0.1% Triton X-100 (Merck, #T8787) for 5 minutes on ice and then centrifuged at 1,300g for 4 minutes. At this step, the supernatant corresponds to the Cytoplasmic Fraction. The pellet was washed in buffer A followed by a centrifugation at 1,300g for 4 minutes. The pellet was resuspended in buffer B (3 mM EDTA, 0.2 mM EGTA, 1mM DTT, 10 mM NaF, 1X Protease inhibitors, 1X Phosphatase inhibitors), kept for 10 minutes on ice and centrifuged at 1,700g for 4 minutes. At this step, the supernatant corresponds to the Soluble Nuclear Fraction. The pellet was resuspended in buffer C (3mM EDTA, 0.2 mM EGTA) and centrifuged at 10,000g for 1 minute. The pellet, corresponding to the Chromatin Fraction, was incubated in 40µL of 1X Blue Loading Buffer (NEB, #B7703S) supplemented with 5 mM MgCl2 and 6µL Benzonase (25units/µL, Merck, #70664) for 1 hour at room temperature. The extracted proteins were then denatured at 95°C for 5 minutes. The α-tubulin and H4 proteins were used as controls for cytoplasmic and chromatin fractions, respectively. Three independent experiments were conducted in **Figures 3G** and **S3B**.

### Western blot

Proteins were separated on 10- or 15-well NuPAGE 4-12% Bis-Tris gels (Thermo Fisher Scientific, #NP0336BOX) and transferred onto nitrocellulose membranes (Thermo Fisher Scientific, #IB23001). Membranes were blocked in TBS-T (Tris 20mM, NaCl 150 mM, Tween® 20 0.05%) with 5% BSA or milk, depending on the used antibody. Primary antibodies were diluted at the appropriate concentration in TBS-T. Secondary antibodies were used as recommended in the manufacturer’s instructions. Blots were imaged with SuperSignal West Femto Chemoluminescence Kit (Thermo Scientific, Cat. No. 34080) and ChemiDoc Imaging System. Relative quantifications were performed with ImageJ software (v1.54g).

### Immunofluorescence with K562 cells

#### Preparation of slides

Approximately 300,000 c/mL asynchronous K562 cells were grown on poly-L-lysine-coated microscope coverslips in a 6-well plate the day before the experiment to attach the suspension cells to the coverslip. Cells were treated during 6 hours with DMSO, 10µM Olaparib or 50µM Veliparib, or during 1 hour with 2mM HU or 100µM H2O2. S-phase cells were labelled by adding 10 µM EdU to the culture media for 2 hours before the end of the drug treatment. Cells were fixed with 4% formaldehyde (Merck, #F8775) for 15 minutes at room temperature (RT) and rinsed twice for 5 minutes with 1X dPBS. Cells were permeabilized with 0.5% Triton-X-100 (Merck, #T8787)/dPBS for 10 minutes at RT. Click-it reactions were performed using the Click-iT® EdU Alexa Fluor® 488 Imaging Kit (Thermo Fisher Scientific, #C10337) according to the manufacturer’s recommendations. Blocking was performed with dPBS containing 0.1% Tween® 20 and 1% BSA for 1 hour at RT. Primary antibodies were diluted in the blocking solution at the appropriate concentration and incubation was performed overnight at 4°C. Cells were washed twice for 5 minutes with 0.05% Tween® 20/dPBS. Secondary antibodies were diluted in the blocking solution at the appropriate concentration and incubation was performed for 90 minutes at RT protected from light. Cells were washed twice for 5 minutes with 0.05% Tween® 20/dPBS and once with dH_2_O. The coverslips were mounted on glass slides with FlouroMount G (SouthernBiotech, #0100-01) and allowed to dry for half a day at RT.

#### Image acquisition and analysis

Fluorescence images were acquired using a confocal LSM 700 run by the Axio Imager software using a 40X/1.3NA oil immersion objective at room temperature. Typically, three images per condition were collected to obtain a total of around 100 cells. Between 10 to 15 Z-stacks per image were collected. From the obtained .czi files, the total number of foci were measured using the SPOT tool in the Imaris (v10.1.1) software. The estimated XY and Z diameters used were 0.5µM and 1µM, respectively. The quality filter was used and adjusted on the control HU condition for each of the studied proteins. For visualization of immunofluorescence images in the figures, all images were processed in ImageJ. Each Z-stack image was reconstructed by maximum intensity Z-axis projection. NB: Wheat Germ Agglutinin (WGA) labels glycoproteins. In permeabilized cells, WGA stains plasma membranes and intracellular compartments such as nucleus or Golgi structures. The number of independent replicates is indicated in figure legends.

### Antibodies used in western blot or immunofluorescence experiments

| Reagent | Source | Used concentration |
| --- | --- | --- |
| Rabbit monoclonal anti-phospho-Histone H2AX (Ser 139) (20E3) | Cell signaling (#9718) | 1:500 (IF) |
| WGA, Alexa Fluor 633 conjugate | Thermo Fisher Scientific (#W21404) | 1:200 (IF) |
| Rabbit monoclonal anti-phospho-CHK1 (Ser345) (133D3) | Cell signaling (#2348) | 1:1000 (WB) |
| Mouse monoclonal anti-CHK1 (2G1D5) | Cell signaling (#2360) | 1:1000 (WB) |
| Mouse monoclonal anti-GAPDH (GA1R) | Thermo Fisher Scientific (#MA5-15738) | 1:2000 (WB) |
| Rabbit monoclonal anti-Poly/Mono-ADP Ribose (E6F6A) | Cell signaling (#83732) | 1:10 000 (WB) |
| Rabbit polyclonal anti-PARP1 | Thermo Fisher Scientific (#PA5-34803) | 1: 10 000 (WB) |
| Mouse anti-αTubulin | Merck (#T9026) | 1:2000 (WB) |
| Rabbit polyclonal anti-ORC1 | Abcam (#ab85830) | 1:2000 (WB) |
| Rabbit monoclonal anti-CDT1 (D10F11) | Cell signaling (#8064) | 1:1000 (WB) |
| Rabbit polyclonal anti-MCM2 | Bethyl (#A300-191A) | 1:10 000 (WB) |
| Rabbit polyclonal anti-phospho-MCM2 (Ser 53) | Bethyl (#A300-756A) | 1:5000 (WB) |
| Mouse polyclonal anti-CDC7 | Cell signaling (#3603) | 1:1000 (WB) |
| Rabbit monoclonal anti-DBF4 (EPR5635) | Abcam (#ab124707) | 1:20 000 |
| Mouse monoclonal anti-CDC45 | Cell signaling (#11881) | 1:1000 (WB) |
| Mouse monoclonal anti-Histone H4 (L64C1) | Cell signaling (#2935) | 1:1000 (WB) |
| Rabbit monoclonal anti-PCNA (D3H8P) | Cell signaling (#13110) | 1:1000 (WB) |
| Mouse monoclonal anti-FANCD2 (FI17) | Santa Cruz Biotechnology (#sc-20022) | 1:250 (WB) |
| Rabbit monoclonal anti-FANCD2 (EPR2302) | Abcam (#ab108928) | 1:500 (IF) |

### DNA fiber assay

DNA fiber assay was performed as previously described [49–51] with minor modifications. Briefly, exponentially growing asynchronous K562 cells were treated during 6 hours with DMSO, Olaparib or Veliparib. Cells were then incubated sequentially with two consecutive thymidine analogs, IdU at 25 µM (Sigma Aldrich, #I7125) and CldU at 250 µM (Sigma Aldrich, #C6891) for 20 minutes each to label in vivo progressing replication forks. Cells were collected and washed with cold dPBS and resuspended at a concentration of 7.5 × 10^5^ cells/mL. A 2 mL drop of cells was put on a positively charged glass slide, allowed to dry for 5 minutes 30 seconds, mixed gently with 7 mL of spreading buffer (200 mM Tris-HCl pH 7.5, 50 mM EDTA, 0.5% SDS) and allowed to settle on the glass slide for 6 minutes 30 seconds. The slide was then tilted at 23° angle to allow the drop to slowly spread. The spread DNA molecules were then air-dried, fixed in methanol/acetic acid and stored at 4°C.

The glass slides were washed in water, denatured in 2.5M hydrochloric acid for an hour, washed with dPBS and washed and incubated for an hour with a blocking solution (dPBS, 0.1% Tween-20, 1% BSA). The immunostaining of IdU and CldU tracks was performed for 45 minutes in a humid 37°C chamber with a mouse anti-BrdU antibody (1:20, BD-Biosciences, #347580) and a rat anti-BrdU antibody (1:100, Abcam, #ab6326), respectively. The slides were then washed with dPBS and 0.1% Tween-20. Secondary antibodies (1:100, anti-mouse Alexa Fluor 568 Invitrogen #A-11004 and 1:100, anti-rat Alexa Fluor 488 Invitrogen #A-21470) were incubated for 30 minutes in a 37°C humid chamber to allow the detection of IdU and CldU, respectively. To ensure that the DNA fibers were not broken during the spreading, DNA was successively immunostained, as a control, with a mouse anti-ssDNA antibody (1:200, Millipore, #MAB3034) followed by an anti-mouse IgG2 Alexa Fluor 647 (1:200, Invitrogen, #A21241). After being washed with dPBS and 0.1% Tween-20, the slides were then air-dried and mounted with FlouroMount G (SouthernBiotech, #0100-01). Fluorescence images of IdU, CldU and DNA control tracks were taken using 63X objective with a Zeiss Axioimager Apotome. The fluorescent tracks were measured using ImageJ software (v1.54g). The length of each labeled DNA track was recorded in µm and converted in kb by using the commonly used conversion factor of 2.59 µm/kb [Jackson and Pombo 1998]. Three independent experiments were conducted.

### iPOND coupled with LC-MS/MS Analysis

#### iPOND

iPOND was performed largely as previously described [52–55] with minor modifications. Briefly, 180×10^6^ K562 cells were pulse labeled with 10 μM EdU for 20 minutes and chase were performed with 10 μM thymidine for 90 minutes. Cells were fixed with 1% formaldehyde (Merck, #F8775) for 15 minutes at RT followed by quenching of formaldehyde by 5 minutes incubation with 0.125 M glycine. Fixed samples were collected by centrifugation at 200g for 10 minutes, washed three times with cold 1X dPBS and stored at −80°C. Cells were permeabilized with 0.5% Triton-X-100 (Merck, #T8787)/dPBS for 30 minutes at RT, washed with 0.5% BSA (Euromedex, #04-100-812)/dPBS and then dPBS. Click chemistry was used to conjugate biotin-TEG-azide (EurogenTech) to EdU-labeled nascent DNA. Click-it reaction was performed for 2 hours at RT protected from light in dPBS containing 10 mM sodium-L-ascorbate, 10 μM biotin-TEG-azide, and 2 mM CuSO4. Cells were re-suspended in lysis buffer (10 mM Hepes-NaOH pH8; 100 mM NaCl; 2 mM EDTA pH8; 1 mM EGTA pH8; 1 mM PMSF; 0.2% SDS; 0.1% Sarkozyl) supplemented with protease (1X cOmplete™ Protease Inhibitor Cocktail, Merck, #11836145001) and phosphatase inhibitors (1X Phosphatase Inhibitor Cocktail Set II, Merck, #524625). Sonication was performed using a Bioruptor Plus sonicator (Diagenode) in 15mL tubes with a metallic bar to reflect the ultrasound within the sample, in a refrigerated water bath, and with 20 cycles of 30 s ON and 30 s OFF at high power. Lysates were centrifuged at 13,200 rpm (HITACHI KOKI Himac CT15E Centrifuge) for 10 minutes at RT. Supernatants were normalized by DNA quantification using a nanodrop device in order to perform iPOND on the same amount of protein in each condition. At this step, a subset of the supernatant was retained as the input. Biotin conjugated DNA–protein complexes were captured using overnight incubation (about 16 hours) at RT with magnetic beads coated with streptavidin (Ademtech, #03213). Captured complexes were sequentially washed with lysis buffer, 500 mM NaCl, 100mM NaCl and dH_2_O. For western blot analyzes of proteins associated with nascent DNA, the beads with DNA-protein complexes attached were resuspended in 1X Laemmli buffer (Merck, #S3401) and eluted by boiling for 30 minutes at 95°C and shaking at 900 rpm. Supernatants were collected and a subset was loaded onto NuPAGE 4-12% Bis-Tris gels (see Western blot section). For LC-MS/MS analyzes of proteins associated with nascent DNA, the beads with DNA-protein complexes attached were resuspended in a small volume of dH_2_O and processed as follows. Six independent experiments were conducted.

#### Material for LC-MS/MS

MS grade Acetonitrile (ACN), MS grade H_2_O and MS grade formic acid (FA) were from ThermoFisher Scientific (Waltham, MA, USA). Sequencing-grade trypsin was from Promega (Madison, WI, USA). Trifluoroacetic acid (TFA), and ammonium bicarbonate (NH4HCO3) were from Sigma-Aldrich (Saint-Louis, MO, USA).

#### Samples preparation prior to LC-MS/MS analysis

Beads from pulldown experiments were incubated overnight at 37°C with 20 μL of 50 mM NH_4_HCO_3_ buffer containing 1 µg of sequencing-grade trypsin. The digested peptides were loaded and desalted on evotips provided by Evosep (Odense, Denmark) according to manufacturer’s procedure before LC-MS/MS analysis.

#### LC-MS/MS acquisition

Samples were analyzed on a timsTOF Pro 2 mass spectrometer (Bruker Daltonics, Bremen, Germany) coupled to an Evosep one system (Evosep, Odense, Denmark) operating with the 30SPD method developed by the manufacturer. Briefly, the method is based on a 44-min gradient and a total cycle time of 48 min with a C18 analytical column (0.15 × 150 mm, 1.9μm beads, #EV-1106) equilibrated at 40°C and operated at a flow rate of 500 nl/min. H_2_O/0.1% FA was used as solvent A and ACN/0.1% FA as solvent B.

The timsTOF Pro 2 was operated in PASEF mode [56] over a 1.3-s cycle time. Mass spectra for MS and MS/MS scans were recorded between 100 and 1700 *m/z*. Ion mobility was set to 0.75-1.25 V·s/cm2 over a ramp time of 180 ms. Data-dependent acquisition was performed using 6 PASEF MS/MS scans per cycle with a near 100% duty cycle. Low *m/z* and singly charged ions were excluded from PASEF precursor selection by applying a filter in the *m/z* and ion mobility space. The dynamic exclusion was activated and set to 0.8 min, and a target value of 16,000 was specified with an intensity threshold of 1000. Collisional energy was ramped stepwise as a function of ion mobility.

#### Data Analysis

MS raw files were processed using PEAKS Online X (build 1.8, Bioinformatics Solutions Inc.). Data were searched against the Homo Sapiens SwissProt database (01_2023, total entries 20,405). Parent mass tolerance was set to 20 parts per million (ppm), with fragment mass tolerance at 0.05 Da. Specific tryptic cleavage was selected and a maximum of two missed cleavages was authorized. For identification, the following posttranslational modifications were included: Acetylation (Protein N-term), oxidation (M), deamidation (NQ), ubiquitin and phosphorylation (STY) as variables and carbamidomethylation (C) as fixed. Identifications were filtered based on a 1% FDR (False Discovery Rate) threshold at both peptide and protein group levels. Label-free quantification was performed using the PEAKS Online X quantification module, allowing a mass tolerance of 10 ppm, a collision cross section (CCS) error tolerance of 0.02 and 0.5min retention time shift tolerance for match between runs. Protein abundance was inferred using the top N peptide method and Total Ion Chromatogram (TIC) was used for normalization. Multivariate statistics on proteins were performed using Qlucore Omics Explorer 3.8 (Qlucore AB, Lund, Sweden). A positive threshold value of 1 was specified to enable a log2 transformation of abundance data for normalization, *i.e.*, all abundance data values below the threshold will be replaced by 1 before the transformation. The transformed data were finally used for statistical analysis, *i.e.*, evaluation of differentially present proteins between two groups using a Student’s bilateral t-test. A p-value better than 0.05 was used to filter differential candidates.

The identification of the 1857 proteins that bind non-specifically to streptavidin beads was performed as follows: from the identification protein file, the average of the six area values was calculated for each identified protein and for each of the pulse conditions (DMSO, Olaparib, Veliparib). From these averages, the Area(drug)/Area(Ctrl click-it neg) ratio was then calculated for each identified protein. If this ratio was ≤ 2, the protein was considered non-specific.

The mass spectrometry proteomics data have been deposited to the ProteomeXchange Consortium via the PRIDE [57] partner repository with the dataset identifier PXD060870.

### Alkaline and Neutral Comet Assay

K562 cells were treated as indicated (**Figures 6A, 6G**). At the end of the treatment, cells were washed twice in cold dPBS. Comet slides were prepared with the CometAssay® Silver Kit (Bio-techne, #4250-050-K). Briefly, cells were resuspended in dPBS at 300 cells/µL and about 8µL of cells + 70µL of low melting agarose were spread in the CometAssay® slide areas. Slides were immersed in lysis solution at 4°C overnight protected from light. Slides were equilibrated in alkaline buffer (300mM NaOH, 1mM EDTA pH >13) for 1 hour or in neutral buffer (1X TBE, Tris-Boric Acid-EDTA pH8) for 15 minutes at 4°C and protected from light. Electrophoresis was performed in alkaline or neutral buffer at 1V/cm for 30 minutes at 4°C and protected from light. Slides were quickly washed twice in dH_2_O for 5 minutes, immersed in 70% ethanol for 30 minutes at RT, protected from light and then allowed to dry for half a day at RT. Comets were stained with 1X SyberGold (Thermo Fisher Scientific, #S11494) diluted in 1X TBE for 20 minutes at RT and protected from light, washed twice for 5 minutes with dH_2_O and allowed to dry for about 1 hour. Fluorescence images were taken using an epifluorescence microscope with a 20X objective and a GFP filter. Tail moment was measured using CometScore software (v2.0.0.38). Two independent experiments were conducted.

### EdU Alkaline and Neutral Comet Assay

K562 cells were treated as indicated (**Figure 6D**). Cells were incubated with 10 µM EdU in culture media for the last 120 minutes of the drug treatment to label S phase cells. Comet slides were prepared as described above. After ethanol incubation, slides were allowed to dry entirely and then, the immunostaining of EdU was performed as follows. The Comet slides were washed with dPBS and 3% BSA for 10 minutes at RT and protected from light. Click-iT reactions were performed using the Click-iT® EdU Alexa Fluor® 647 Imaging Kit (Thermo Fisher Scientific, #C10337) according to the manufacturer’s recommendations. The spots on comet slides were covered each with ∼100µL of Click-iT solution and Click-iT reactions were performed for 30 minutes at RT and protected from light. Comet slides were washed with dPBS and 3% BSA for 10 minutes and then with dPBS for 10 minutes, both at RT and protected from light. Comets were stained with SyberGold as described above and allowed to dry entirely at RT, protected from light. Fluorescence images were taken using 20X objective with a Zeiss Axioimager Apotome. The GFP and Cy5 channels were used for the measurement of SyberGold and Alexa-fluor 647 signals, respectively. Tail moment of EdU negative comets was measured using CometScore software (v2.0.0.38). Two independent experiments were conducted.

### Replication Timing Analysis

#### RT Analysis

About 15×10^6^ K562 cells were incubated with 50μM BrdU (5-bromo-2’-desoxyuridine) (abcam, #ab142567) during 2 hours before the ending point of treatments, harvested and fixed with 30% dPBS 1X and 70% cold ethanol. Fixed cells were treated with RNaseA (0,5mg/ml) (Merck, #10109169001) and propidium iodide (50μg/ml) (Thermo Fisher Scientific, #P3566) during 30 minutes at room temperature before cell sorting. 150,000 cells were sorted in two fractions using FACSAria Fusion (BD Biosciences). Cells were incubated in lysis buffer (50mM Tris pH8; 10mM EDTA, 300mM NaCl, 0.5% SDS) and proteinase K (Thermo Fisher Scientific, #EO-0491) (0.2 mg/ml) during 3 hours at 65°C. DNA was extracted using phenol-chloroform and precipitated with 2.5 volume of cold 100% ethanol, 2µL glycogen (Merck, #10901393001) and 25% sodium acetate (3M, pH=5.5; Invitrogen #AM9740). After precipitation, pellets were rinsed with 75% ethanol, dried and resuspended in 100µL Tris buffer (10mM, pH=8). DNA was sonicated to obtain fragments between 500 and 1,000 bp. After sonication, DNA was denatured at 95°C during 5 minutes and kept on ice for 10 minutes. Immunoprecipitation of nascent DNA was performed as described in our previous study [58]. Briefly, nascent DNA was immunoprecipitated using the IP-STAR apparatus (Diagenode) with BrdU antibody (10μg, Anti-BrdU Pure, BD Bioscience, #347580). Immunoprecipitated nascent DNA was purified and precipitated as mentioned above. DNA was then amplified using Seqplex as mentioned by the manufacturer (Sigma, #SEQXE). Each fraction of sorted cells is labeled either with Cy3 or Cy5 ULS molecules, respectively (KREATECH, #EA-005) and as recommended by the manufacturer. The hybridization was performed according to the manufacturer instructions on 4×180K human microarrays (GenetiSure Cyto CGH Microarray Kit, 4X180K, AGILENT Technologies, genome reference hg19, G5983A). Feature extraction was performed with the Feature Extraction 9.1 software (Agilent Technologies). Microarrays were scanned with an Agilent’s High-resolution C scanner using a resolution of 3μm and the autofocus option. The START-R suite was used for analysis of replication timing [58]. Differential analysis of two experiments, each composed of two independent biological replicates, were performed with START-R Analyzer and visualized with START-R Viewer.

Data are deposited in GEO with accession number GSE289208.

#### Genomic studies of advanced and delayed RT domains

For each experiment, START-R Analyzer generated segmentation bed files corresponding to early, mid, late, TTR, advanced, and delayed replicating domains. The percentage in GC was obtained with the “Extract Genomic DNA using coordinates from assembled/unassembled genomes” (Galaxy Version 3.0.3+galaxy3) and the “geecee” (Galaxy Version 5.0.0) softwares. The coverage of the different replicating domains with large genes (>400 kb), constitutive origins, CpG islands, and putative G4 were done with the “Coverage of a set of intervals on second set of intervals” software (Galaxy Version 1.0.0). Boxplots illustrating differences in these coverages or percentages were generated with RStudio (version 2024.12.0). The positions of K562 replication origins and putative G4 (hg19 assembly) were taken from Picard *et al.* (2014) [59]. The positions of genes came from the UCSC table browser RefSeq Genes database without duplicates (hg19 assembly). Large gene regions (>400 kb) were extracted from the aforementioned gene database. The positions of CpG islands came from the UCSC table browser (hg19 assembly). The positions of CFS (hg38 assembly) were taken from the HumCFS database [60] and converted with LiftOver to hg19 genome assembly. The “Join the intervals of two datasets side-by-side” software (Galaxy Version 1.0.0) was used for the identification of CFS in the 54 regions where RT is delayed by veliparib in **Supplemental table 3**. For the identification of genes in the regions where RT is specifically delayed by veliparib (**Figure 7G** and **Supplemental table 4**), the positions of genes were imported from the UCSC table browser RefSeq Genes database (hg19 assembly). Then, the “Intersect the intervals of two datasets” software (Galaxy Version 1.0.0) was used to list the genes located in the 51 genomic regions where RT is specifically delayed by veliparib. Finally, the “annoteMyIDs” software (Galaxy Version 3.18.0+galaxy0) was used to convert RefSeq ID in Gene Symbol and Gene Name.

### Gene Ontology analysis

Gene Ontology analysis was performed with the ToppFun tool from the ToppGene Suite (https://toppgene.cchmc.org/), using the May 2023 version for **Figure S4C** and the January 2025 version for **Figures 4D** and **7G**. The chosen pValue method was the Probability density function, with the Bonferroni correction and a 0.05 p-Value cutoff.

### Statistical analysis

Statistical analyses, boxplots, barplots and volcanoplots were obtained using R 4.3.3 environment and ggplot2, ggsignif and ggrepel R packages.

## RESULTS

### Inhibition of PARP activity directly delays the S phase progression and induces replicative stress

To investigate the immediate effect of an absence of PARylation within a single S phase in cultured cells, we used competitive inhibitors of PARP1 and PARP2. Two PARP inhibitors, namely olaparib and veliparib, were used to target the catalytic activity of PARP1 and PARP2 and quickly induce an absence of PARylation. To precisely analyze the relationship between PARylation and DNA replication within a single S phase and to avoid cross-effects between successive cell cycles, we studied the effect of PARP inhibitors on synchronized K562 human cells. While avoiding the use of drugs to synchronize cells, the elutriation technique allows us to obtain synchronous cell populations highly enriched for each phase of the cell cycle. Cells were synchronized in early G1 and cultured with olaparib or veliparib until the end of the S phase (**Figure 1A**). The progression of cells in S phase was measured at different timepoints by flow cytometry analyses (**Figure 1A**). We found that both inhibitors induce an accumulation of cells in S phase during the first cell cycle following cell synchronization and treatment (**Figures 1A, 1B**). Importantly, cells exposed to these inhibitors does not stay in G1 (**Figures 1A and S1A**) indicating that absence of PARP1-2 activity does not affect the entry in S phase. According to the cell cycle profiles (**Figure 1A**), cells in the control condition enter G2 14 hours after synchronization in early G1, whereas cells in the PARPi condition enter G2 24 hours after the synchronization, indicating that PARPi cause a delay in S phase progression of about 10 hours. These observations strongly suggest that an absence of PARylation causes a direct delay of S phase progression due to events occurring during this S phase, rather than in a previous S phase, as previously described with the induction of post-replication gaps [33]. An analysis of the cell cycle progression of G2-synchronized WT K562 cells enabled us to estimate that the approximate duration of the G1 and S phases are 8 and 10 hours, respectively, in this cell line (**Figure S1B**). Thus, we assumed that the effect of PARP inhibitors during a single S phase can be studied by short treatments (from 2 to 10 hours) of exponentially growing asynchronous K562 cells.

**Figure 1:**
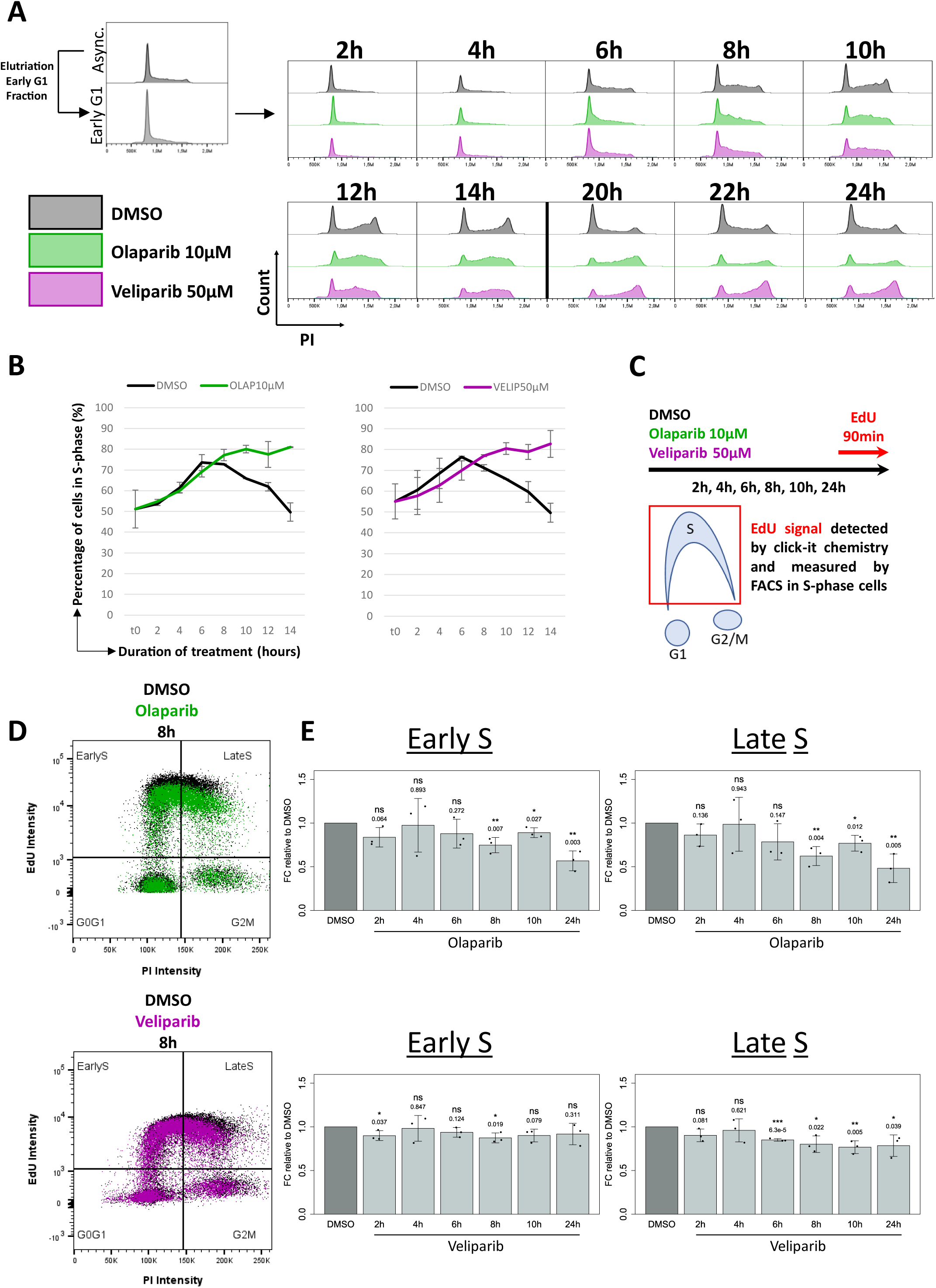
Absence of PARylation impairs DNA replication during the same S phase. **(A)** Cell cycle progression of K562 cells synchronized in early G1 by elutriation. Cells were treated with olaparib 10µM (green) or veliparib 50µM (purple) from early G1 to the indicated timepoints. **(B)** Quantification of cells in S phase. SD are indicated (n=2). **(C)** Schematic of the experiment for EdU incorporation in exponentially growing asynchronous K562 cells treated with olaparib or veliparib for the indicated periods. **(D)** Cell cycle profiles after EdU incorporation. An example is shown for the 8h timepoint. **(E)** Histograms representing the relative signal of EdU in early or late S phase in K562 cells normalized to the control. SD are indicated (n=3). Statistics: for each timepoint, unpaired Student’s t-test was performed to compare the DMSO versus PARPi conditions. ns=non-significant, * p<0.05, ** p < 0.01, *** p < 0.001.

We then analyzed the impact of an absence of PARylation on DNA synthesis during a single S phase. Asynchronous exponentially growing K562 cells were exposed to PARP inhibitors for 2 to 10 hours, and the incorporation of the thymidine analog 5-ethynyl-2′-deoxyuridine (EdU) was measured in S phase cells (**Figure 1C**). A 24-hour time point was also included for comparison with published results [33,37]. We found that olaparib and veliparib induce a progressive decrease in DNA synthesis throughout the time course of treatment (**Figures 1D, 1E**). Reduced EdU incorporation in S phase cells is caused by either reduced origin firing or decreased or even stopped replication fork progression. The same effect was observed for the two inhibitors when the experiment was performed in RPE1 cells (**Figures S1C, S1D**). In both cell lines, decrease in DNA synthesis was slightly higher in late *versus* early S phase cells (**Figures 1D, 1E, S1C and S1D**).

The effect of a lack of PARylation on ATR checkpoint activation was then analyzed by total protein extraction and western blot in the same experimental design. We showed that PARP inhibitors rapidly induce a mild increase in the level of CHK1 phosphorylation at residue S345 indicating an increase in replication stress (**Figures 2A, 2B**). Another ATR-downstream target, namely the phosphorylation of H2AX at residue S139 (γH2AX) was also analyzed. Thus, immunofluorescence was performed on asynchronous cells treated for 6 hours with olaparib or veliparib and γH2AX foci were enumerated in S phase cells visible by EdU staining (**Figure 2C**). We observed an increase in the number of γH2AX foci both in early or mid-late S phase cells after a short treatment with PARP inhibitors (**Figures 2D, 2E**), indicating either DNA breaks formation or induction of replication stress as previously published [61]. The induction of γH2AX foci by PARP inhibitors was specific to S-phase cells, as indicated by the comparison between EdU negative cells (**Figure S2A**) *versus* EdU positive cells (**Figure 2E**). In addition, this increase is slightly more pronounced in mid-late *versus* early S phase cells (**Figure 2E right vs left**). Thus, an absence of PARylation during a single S phase induces replication stress.

**Figure 2:**
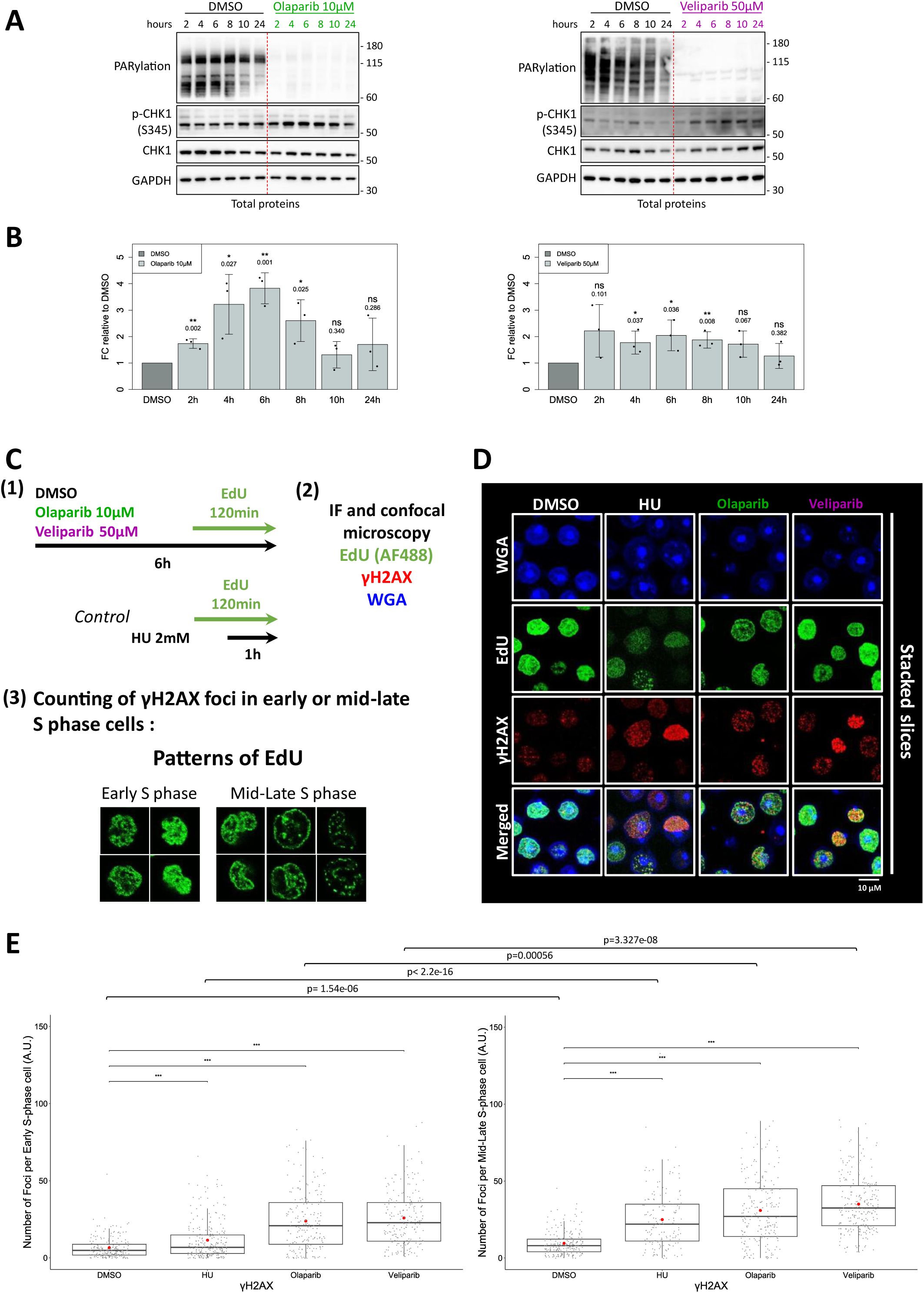
Absence of PARylation induces replicative stress during the same S phase. **(A)** Total protein levels of Chk1, Chk1 phosphorylated on Serine 345 residue and PARylated proteins were analyzed by western blot after increasing treatment times with olaparib 10µM or veliparib 50µM in exponentially growing asynchronous K562 cells. GAPDH was used as loading control. **(B)** Histograms representing the relative amount of phosphorylated CHK1 normalized to the control. SD are indicated (n=3). Statistics: for each timepoint, unpaired Student’s t-test was performed to compare the DMSO versus PARPi conditions. ns=non-significant, * p<0.05, ** p < 0.01, *** p < 0.001. **(C)** Schematic of the experiment. Exponentially growing asynchronous K562 cells were treated with olaparib 10µM or veliparib 50µM for 6 hours. A treatment with HU 2mM for 1 hour was also performed as a positive control for DNA damage and replicative stress induction. S phase cells are labelled by EdU. Early S phase cells have a fuzzy EdU signal throughout the nucleus, while mid-late S phase have a more punctuated EdU signal pattern located at the side of the nucleus. Wheat Germ Agglutinin (WGA) allows the detection of cellular and nuclear membranes. **(D)** Immunofluorescence of γH2AX, EdU and WGA signals. **(E)** Boxplots indicating quantification of γH2AX foci per Early or Mid-Late S-phase cells. A total of 200 to 250 cells from 3 independent experiments were quantified for each condition. The mean values are indicated in red on boxplots (DMSO=6.58; HU=11.44; Olaparib=23.90; Veliparib=25.86 in early S phase cells and DMSO=9.54; HU=24.93; Olaparib=30.91; Veliparib=35.00 in mid-late S phase cells). Statistics: non-parametric Mann-Whitney test. ns=non-significant, * p<0.05, ** p < 0.01, *** p < 0.001.

### Olaparib and veliparib differentially impact fork progression and origin firing

The olaparib and veliparib inhibitors have already been demonstrated to accelerate replication fork speed without affecting fork symmetry with long lasting treatments [37,38]. To test the effect of these PARP inhibitors on replication fork speed and fork symmetry for a shorter duration, we performed DNA fiber assays using sequential, 20-minute pulses of the thymidine analogs IdU and CldU (**Figure 3A**). Single-strand DNA staining was also performed to control the DNA fiber integrity. Unexpectedly, only olaparib mildly increased the speed of fork progression by approximately 12% in K562 cells (**Figure 3B**). In contrast, only veliparib affected fork symmetry (**Figure 3C**) indicating that this inhibitor induces more fork stalling and/or collapse than olaparib.

**Figure 3:**
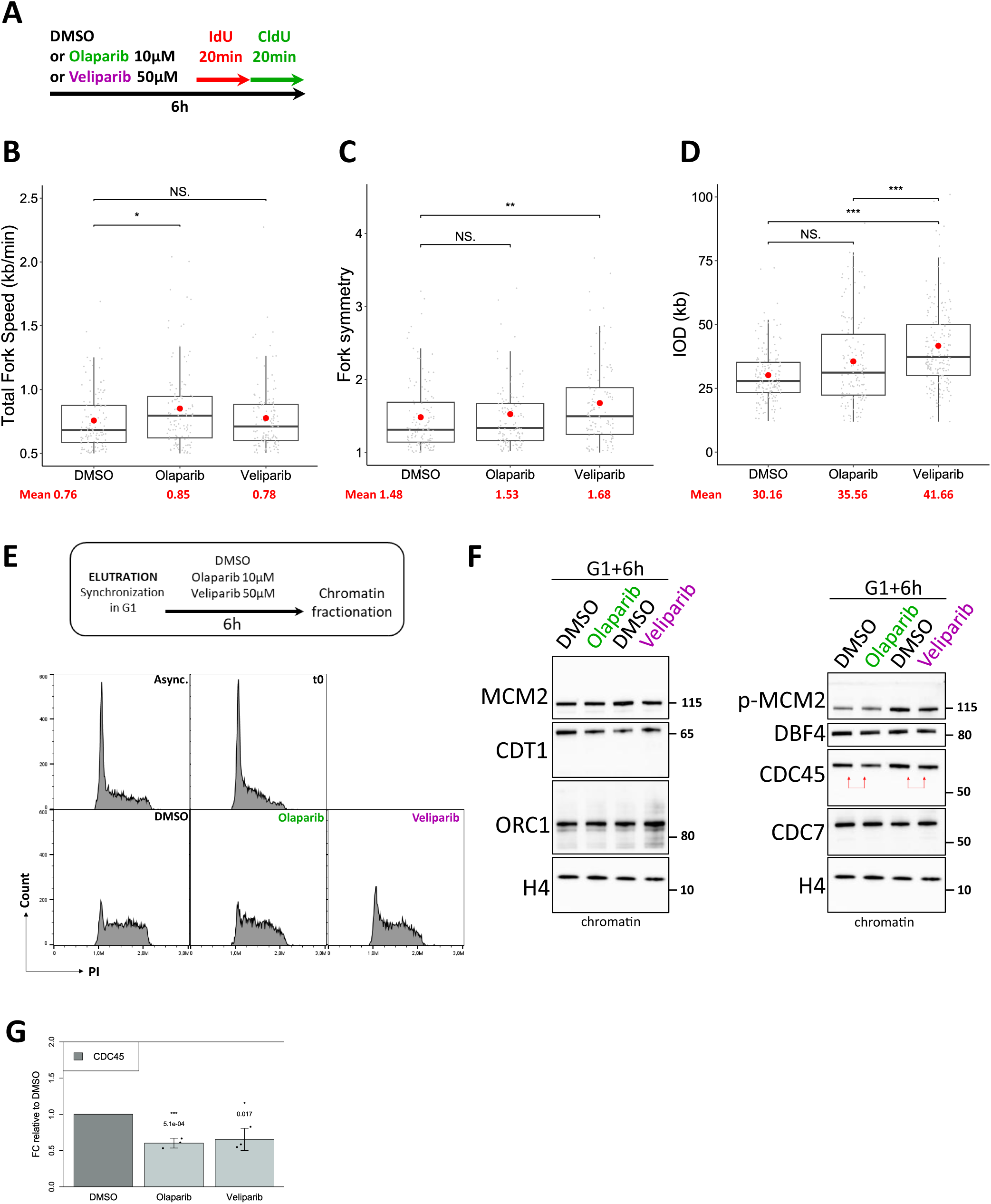
Absence of PARylation decreases origin firing during the same S phase. **(A)** Schematic of the DNA fiber assay. Exponentially growing asynchronous K562 cells were treated with olaparib 10µM or veliparib 50µM for 6 hours. IdU (red) was added for 20 min followed by CldU (green) for 20 min, before the end of the treatment. **(B)** Boxplot representing the fork speed (kb/min) in each condition. 119 forks were scored from 3 independent experiments in each condition. **(C)** Boxplot representing the IdU/CldU or the CldU/IdU ratio ≥1 in each condition from values in **(B)**. **(D)** Boxplot representing the inter-origin distance (IOD) (kb) in each condition. Scored IOD from 3 independent experiments: control (DMSO) = 164; olaparib = 142; veliparib = 190. **(B-D)** Mean is indicated in red. Statistics: non-parametric Mann-Whitney test. ns=non-significant, * p<0.05, ** p < 0.01, *** p < 0.001. **(E)** Schematic of the chromatin fractionation assay. K562 cells were synchronized in G1 and treated with olaparib 10µM or veliparib 50µM for 6 hours until the mid S phase. At the end of the treatment, chromatin proteins were extracted. Cell cycle profiles after synchronization by elutriation are shown. **(F)** Western blot of pre-replication complex (preRC) proteins (MCM2, CDT1, ORC1) and pre-initiation complex (preIC) proteins (phosphorylated MCM2, DBF4, CDC45, CDC7) in chromatin protein extracts. H4 was used as loading control. Red arrows highlight the slight decrease of chromatin-bound CDC45 in treated cells. **(G)** Histogram representing the relative amount of the CDC45 protein normalized to the control. SD are indicated (n=3). Statistics: unpaired Student’s t-test was performed to compare the DMSO versus PARPi conditions. ns=non-significant, * p<0.05, ** p < 0.01, *** p < 0.001.

We have shown that PARP inhibition reduced the global rate of DNA synthesis (**Figures 1D, 1E**), but increased replication fork speed with olaparib (**Figure 3B**). It has been shown that an increase in fork progression is usually concomitant with a reduction in the number of active origins and *vice versa* [62], although the molecular mechanisms regulating this relationship remain to be elucidated. This prompts the question of whether PARP1/2 inhibition impacts the activity of replication origins. In a previous study, olaparib (10µM for 24h) was shown to cause an increase in fork speed correlated with a decrease in origin firing [37]. A recent study has shown that this effect on origin firing is a secondary response following the increase in fork speed [39]. In order to investigate the activity of origins when PARylation is suppressed within a single S phase, we analyzed the inter-origin distance (IOD) in K562 cells treated with PARP inhibitors for a short period (**Figure 3A**). We found that olaparib slightly increased the IOD, even if this difference did not reach significance. This indicates a modest decrease in origin firing (**Figure 3D**). This decrease is nevertheless coupled with an increase in fork speed (**Figure 3B**), as previously published. Unexpectedly, the veliparib inhibitor induced a much higher increase in IOD (**Figure 3D**) without altering replication fork speed (**Figure 3B**). This suggests that this inhibitor has a specific and major effect on origin activity. It is important to note that an increase in IOD reflects a reduction in origin firing within clusters, but does not rule out changes (more or less) in the number of clusters that are activated at a given time.

To further understand the molecular effect of an absence of PARylation on the activity of replication origins, we analyzed the formation at chromatin during S phase of the pre-replication complex (preRC) at the replication origin and of the pre-initiation complex (preIC), which is essential for the activation of origins. Extraction of chromatin proteins (Controls in **Figure S3A**) was performed after cell synchronization in G1 and incubation with PARP inhibitors for 6 hours, until the mid S phase (**Figure 3E**). The chromatin abundance of the preRC proteins was not modified by PARP inhibitors (MCM2, CDT1 and ORC1 in **Figures 3F, S3B**). However, a slight, yet significant, chromatin-binding defect of CDC45, a preIC component, was observed after PARP inhibition (**Figures 3F, 3G**). The abundance of CDC45 in a total protein extract was not altered by PARP inhibition under similar conditions (**Figure S3C**). CDC45 is recruited to the preIC after MCM phosphorylation by CDK and DDK protein complexes, allowing the formation of an active helicase CMG (CDC45-MCM-GINS) complex. The DDK complex is composed of the proteins DBF4 and CDC7. It is known that CHK1 phosphorylation by ATR can lead to inactivation of late origins by inhibiting DDK and therefore resulting in a decrease in CDC45 chromatin loading at late origins [for a review, 63]. We have shown that absence of PARylation increases CHK1 phosphorylation when cells are exposed to PARPi during a single S phase (**Figures 2A, 2B**). However, the chromatin abundance of both DBF4 and CDC7 proteins was not altered by PARP inhibition (**Figures 3F, S3B**). Consistently, the phosphorylation of MCM2 was not significantly reduced by PARP inhibitors (**Figure S3B**). These results therefore suggest that the reduction of CDC45 chromatin assembly (**Figures 3F, 3G**) and origin firing (**Figure 3C**) induced by PARP inhibitors are not due to CHK1 activation (**Figure 2**). This could result from another unknown molecular mechanism, maybe induced as a secondary response following the increase in fork speed when PARylation is inhibited, as recently published [39].

### PARP inhibition alters the protein composition of the replisome

As PARylation is known to control protein-protein interactions, we next investigated whether a lack of PARylation impacts the protein composition of the elongation machinery during DNA synthesis. To identify the proteins that are specific to the replication forks, Isolation of Proteins On Nascent DNA (iPOND) was performed as previously described [52–55,64–66]. Briefly, cells are first exposed to a short pulse of EdU to label nascent DNA (**Figure 4A**). Protein-DNA complexes are then maintained by a cross-linking step and labelled with biotin by a click-it reaction. Finally, protein-DNA complexes are extracted by sonication and chromatin purification, and protein-nascent DNA complexes are enriched by immunoprecipitation with streptavidin beads. iPOND was coupled with LC-MS/MS to characterize the whole replisome proteome in K562 cells in physiological conditions or subjected to an absence of PARylation. iPOND-LC-MS/MS was performed on exponentially growing asynchronous K562 cells treated with DMSO, olaparib or veliparib for 6 hours and with EdU during the last 20 minutes (“DMSO pulse”, “Olaparib pulse” or “Veliparib pulse”). We also treated K562 cells with EdU for 20 minutes followed by a 90-minute chase in media without EdU (“DMSO chase”) (**Figure 4A**). By using a 1% false discovery rate (FDR), a total of 4060 individual proteins were identified from six biological independent experiments per condition. The specificity of replication fork protein enrichment (pulse samples) over bulk chromatin (chase sample) was confirmed by PCA analysis (**Figure S4A, left**) and western blot (**Figure S4B**). A technical control consisting of performing iPOND-LC-MS/MS on cells without the click-it reaction (“Ctrl click-it neg”) allowed us to identify 1857 proteins that unspecifically bind streptavidin beads (see materials and methods). We first compared “DMSO pulse” to “DMSO chase” (unilateral t.test, p-value 0.05) in order to identify the replisome proteome of K562 cells under physiological conditions. The result is a ratio of the relative abundance of each protein purified with nascent DNA compared to bulk chromatin. A total of 982 proteins were identified significantly enriched in the “DMSO pulse” condition (fold change > 1). Further filtering of this list by using a unique peptide ≥ 2 cutoff and removing proteins identified in the “Ctrl click-it neg” results in a list of 686 proteins enriched in the K562 replisome under physiological conditions (**Figures 4A, 4B** and **Supplemental Table 1**). The significant enrichment for DNA replication process among the top ten Gene Ontology (GO) terms was confirmed by GO analysis (**Figure S4C**). We also compared our list of 686 proteins with the “catalogue of 593 replication fork proteins in human cells” published by the Cortez’s lab [66]. We found that a total of 49% (290 proteins) are shared with our nascent DNA-associated K562 proteome (**Figure S4D**). Importantly, we detected in control cells that PARP1 and PARG proteins are more enriched in nascent DNA than in chromatin DNA during DNA replication, highlighting a potential role for PARylation at the replisome under physiological conditions (**Figure 4B**). We then examined whether olaparib or veliparib altered the abundance of each of these 686 proteins within the replisome by comparing “DMSO pulse” to “Olaparib pulse” or “Veliparib pulse” (bilateral t.test, p-value 0.05) (**Figures 4A and S4A, right**). Four proteins are found enriched and 13 proteins depleted after olaparib treatment (**Figure 4C, left**) whereas 25 proteins are depleted and no enriched protein after veliparib treatment (**Figure 4C, right**) (**Supplemental Table 2**). Among the depleted proteins, we observed by GO analysis a significant reduction within the replisome of proteins involved in the DNA damage response related to the homologous recombination repair pathway (**Figure 4D**). In particular, the proteins FANCD2, FANCI, BRCA1, BARD1, RNF168, and RAD18 were among the most significantly reduced after PARP inhibition (**Figure 4C**). Importantly, their decrease is not simply due to reduced capture of EdU-labelled DNA, since the same amounts of iPOND-isolated proteins were injected into the mass spectrometer for each condition, and the levels of key replisome factors such as PCNA or DNA polymerases remain relatively unchanged between the different conditions (**Figure S4E**). In addition, since auto-PARylated PARP1 is also able to regulate the expression of some genes [for a review, 2], PARP inhibitors could reduce the gene expression of the protein candidates identified above. However, we found no difference in the protein level of FANCD2 between total protein extracts from untreated and treated cells (**Figures S4F, S4G**), confirming for FANCD2 and suggesting for the other candidates that PARP inhibitors do not alter the expression of these genes. Therefore, these results suggest that PARylation may regulate the recruitment or the stabilization of these proteins at the replisome.

**Figure 4:**
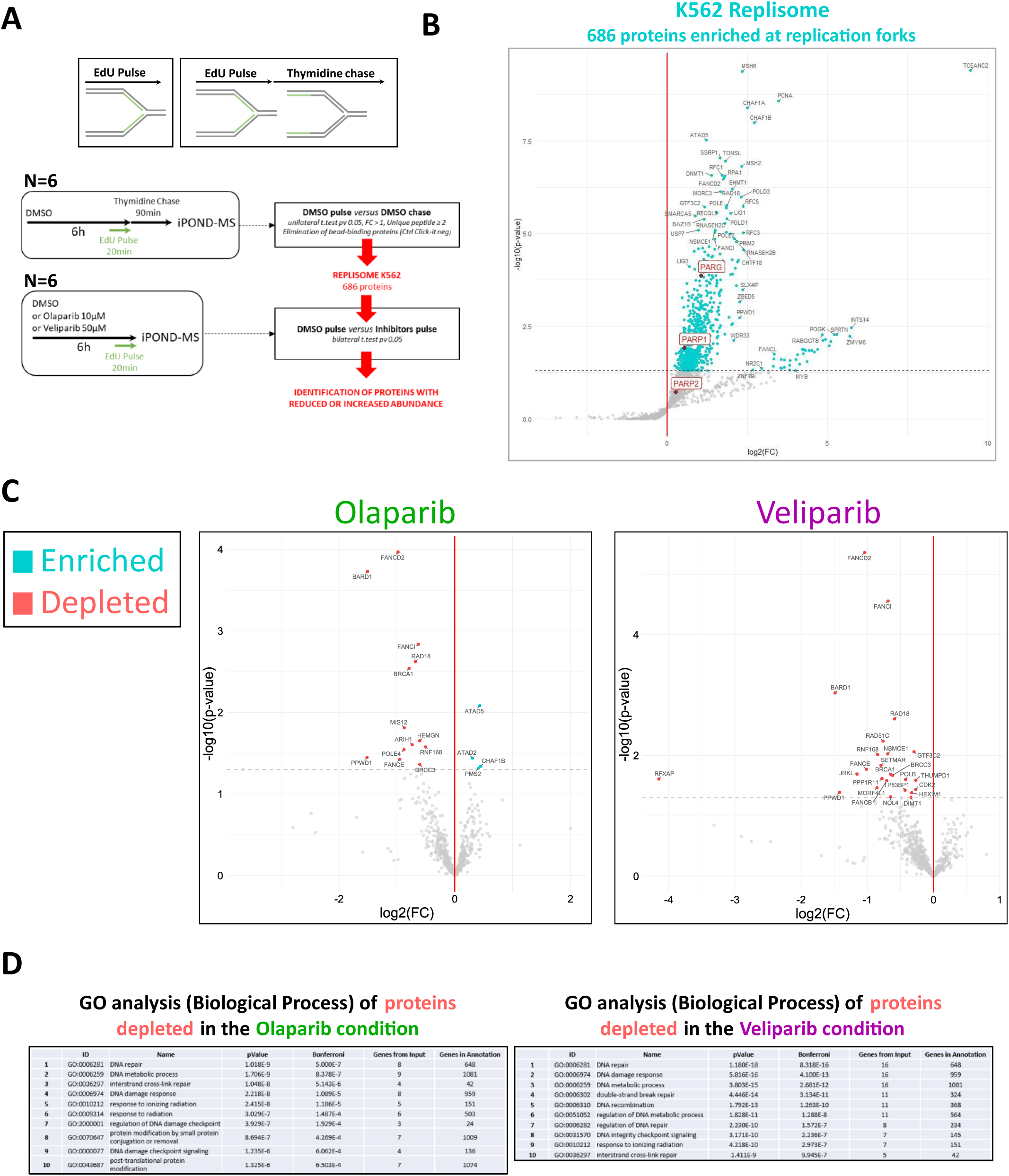
Absence of PARylation modifies the protein composition of the replisome during the same S phase. **(A)** Schematic of the iPOND-MS assay. Exponentially growing asynchronous K562 cells were pulsed for 20 minutes with EdU 10µM and chased for 90 minutes with thymidine 10μM. Comparison between pulse and chase samples allowed to identify the replisome proteome of the K562 cell line. To characterize the replisome proteome upon PARylation dysregulation, K562 cells were treated with olaparib 10µM or veliparib 50µM for 6 hours and incubated with EdU 10µM for the last 20 minutes of the treatment. Protein extracts enriched by iPOND were analyzed by LC-MS/MS. **(B)** Volcano plot indicating the 686 proteins identified as replisome-specific in the K562 cell line. The x-axis and y-axis represent the p-value and fold change, respectively. The dotted line indicates the 0.05 p-value threshold. Statistics: unilateral Student’s t-test with a 0.05 p-value performed between the “DMSO Pulse” and “DMSO chase” datasets (n=6). **(C)** Volcano plots indicating the replisome proteins identified in **(B)** which are enriched (blue) or depleted (red) when K562 cells are exposed to olaparib 10µM or veliparib 50µM for 6 hours. Statistics: bilateral Student’s t-test with a 0.05 p-value performed between the “DMSO Pulse” and “Olaparib Pulse” or “Veliparib Pulse” datasets for each of the 686 proteins identified in **(B)** (n=6). **(D)** Gene Ontology analysis (ToppGene suite) performed on the gene names of the depleted proteins identified in **(C)**.

During DNA replication, the E3 ubiquitin-protein ligase RAD18 is required for monoubiquitination of PCNA on lysine 164 and activation of Translesion DNA synthesis to allow DNA lesion bypass [67,68]. Conversely, the protein ATAD5, found enriched in the olaparib replisome (**Figure 4C**), promotes deubiquitination of PCNA and PCNA unloading from chromatin [69]. We thus investigated the status of chromatin-bound PCNA during a single S phase in the context of PARP inhibition. Extraction of chromatin proteins was performed after cell synchronization in the early-S phase and incubation with olaparib or veliparib for 6 hours, until the mid-late S phase (**Figure S5A, left**). A slight increase in chromatin-bound PCNA was observed in cells treated with PARP inhibitors (**Figure S5A, lanes 1 to 3**). A slight increase in the level of PCNA within replisome was also observed in the iPOND experiments in cells exposed to these drugs (**Figures S4B right blot, lane 7 *vs* lanes 9 and 10**), yet not significant (**Figure S4E**). Conversely, a reduction in the monoubiquitinated form of PCNA was observed in chromatin extracts from cells treated with PARP inhibitors (**Figure S5A, right, lanes 1 to 3**), consistent with the decrease in RAD18 abundance within olaparib and veliparib replisomes (**Figure 4C**). An analysis of differential PTM abundance between the “DMSO pulse” and “Olaparib pulse” or “Veliparib pulse” iPOND samples corroborated this result (**Figure S5B**). These results suggest that a lack of PARylation may inactivate translesion synthesis at the replisome.

Consequently, PARylation seems important to regulate the proteome composition of the replicative machinery during S phase and might regulate some fork restart pathways such as TLS.

### PARP inhibition induces formation of FANCD2 foci and double-strand breaks during S-phase due to an accumulation of single-strand breaks in the DNA template

We then investigated whether an absence of PARylation could reduce the chromatin loading of the protein candidates identified in the iPOND analyses. We focused on the FANCD2 protein and analyzed the foci formation of this protein by immunofluorescence performed on asynchronous cells treated for 6 hours with olaparib or veliparib (**Figure 5A**). We also included treatments with H2O2 or HU for controls. Surprisingly, we found that the two PARPi induced a significant increase in the number of FANCD2 foci in S-phase cells, with olaparib displaying the highest increase (**Figures 5B, 5C**). This increase was much more pronounced in late S-phase cells than in early S-phase cells (**Figure 5C, left *vs* right**). This result could be not very intuitive given the iPOND results (**Figure 4C**). However, if these FANCD2 foci are associated with fork collapse, it is important to note that these events are not associated with DNA synthesis and could not be detected by the iPOND experiment, especially given the experimental design chosen here (*ie.* 20-minute EdU pulse after several hours of PARP inhibition, **Figure 4A**).

**Figure 5:**
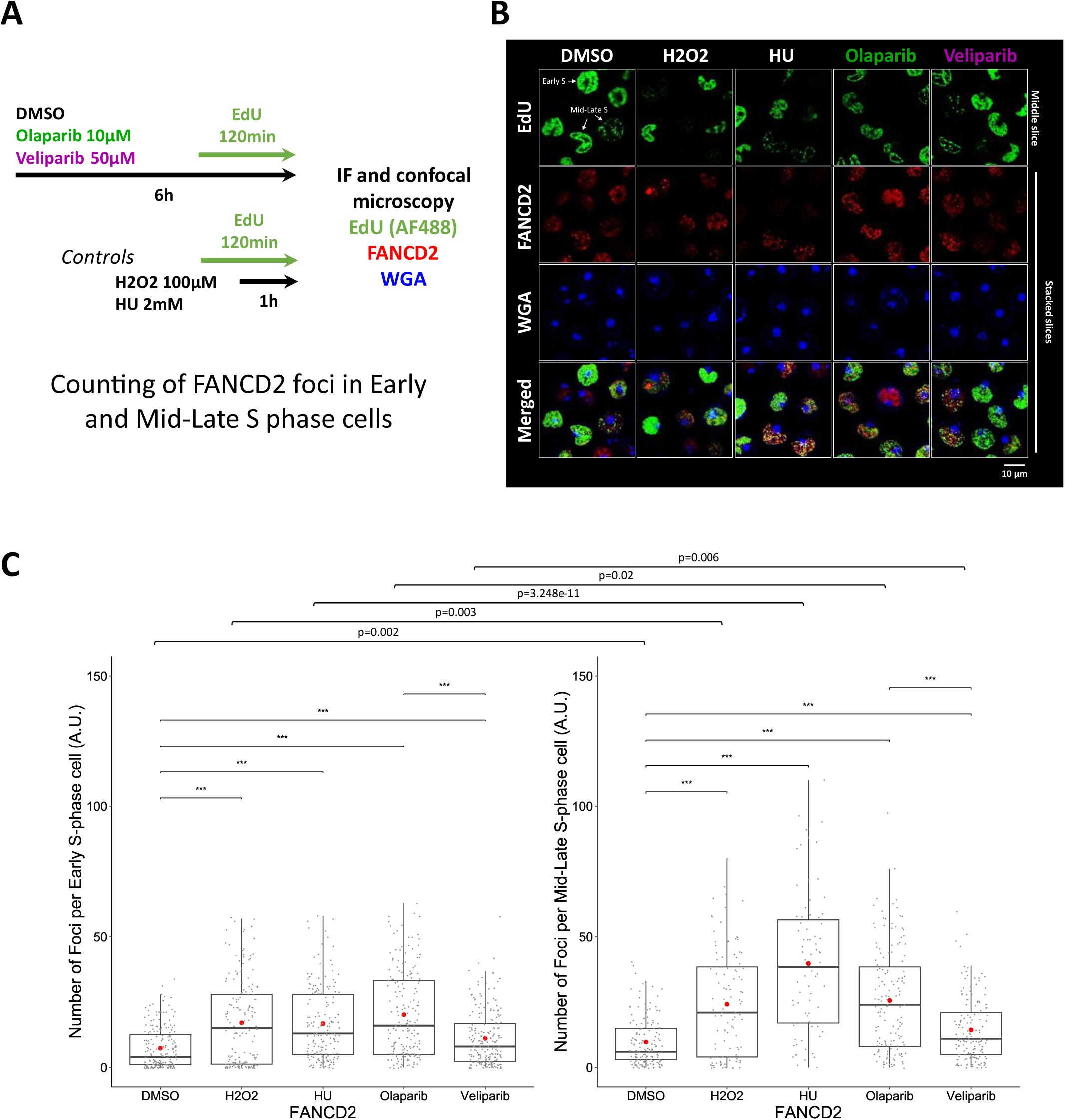
Absence of PARylation induces an increase in FANCD2 foci during the same S phase. **(A)** Schematic of the experiment. Exponentially growing asynchronous K562 cells treated with olaparib 10µM or veliparib 50µM for 6 hours. H2O2 100µM or HU 2mM for 1 hour were also performed as a positive control for DNA damage induction. S phase cells are labelled by EdU. Early S phase cells have a fuzzy EdU signal throughout the nucleus, while mid-late S phase have an EdU signal only at the side of the nucleus. Wheat Germ Agglutinin (WGA) allows the detection of cellular and nuclear membranes. **(B)** Immunofluorescence of FANCD2 signal. **(C)** Boxplots indicating quantification of FANCD2 foci per early or mid-late S-phase cells. A total of 150 to 200 cells from 2 independent experiments were quantified for each condition. The mean values are indicated in red on boxplots (DMSO=7.37; H2O2=17.08; HU=16.77; Olaparib=20.20; Veliparib=11.13 in early S phase cells and DMSO=9.71; H2O2=24.20; HU=39.78; Olaparib=25.64; Veliparib=14.37 in mid-late S phase cells). Statistics: non-parametric Mann-Whitney test. ns=non-significant, * p<0.05, ** p < 0.01, *** p < 0.001.

Furthermore, the FANC-BRCA pathway is known to be involved in the repair of some double-strand breaks (DSB) by homologous recombination (HR) [for two reviews, 70,71]. During S phase, one-ended DSB, a subtype of DSB, can be formed when replication forks encounter SSBs in the DNA template, resulting in fork collapse. During the BER pathway, a nucleotide excision occurs, which results in a transient DNA SSB. These BER intermediates are an important source of SSBs in cells since thousands of endogenous base damages are formed each day in every human cell [for a review 72]. PARP1 activity has a key role in SSBR *via* XCCR1 recruitment, thereby contributing to the repair of SSB formed during BER [for a review 73]. It has recently been shown that unrepaired BER intermediates in template DNA trigger replication fork collapse when cells lack PARP activity [46]. However, to our knowledge, there is no direct evidence that PARP inhibition in physiological conditions leads to the accumulation of endogenous unrepaired SSBs in the template DNA strands. To directly test this hypothesis, we employed comet assays to measure the amounts of breaks in DNA. Comet assay performed under alkaline conditions reveals SSBs, DSBs and also abasic or alkali-labile sites whereas when performed under neutral conditions, it reveals only DSBs. To ensure that we were measuring unrepaired SSBs in the DNA template, and not post-replicative single-strand gaps generated by PARP inhibitors behind the replication forks during S phase, cells were synchronized in G2 and cultured with PARPi until the end of G1 (**Figure 6A**). A treatment with H2O2 was also included as a control for the induction of DNA damages. An increase in tail moment was observed in the alkaline comet assay when cells were exposed to PARP inhibitors during G1, at a level similar to that observed when cells are subjected to H2O2 (**Figure 6B**). Such an increase was not observed in the neutral comet assay (**Figure 6C**), indicating that DNA breaks observed in the alkaline comet assay were mostly SSBs or base damages and not DSBs. Since PARylation is involved in SSBR, we assume that the DNA damages observed in the alkaline comet assay in the PARPi condition are mostly endogenous unrepaired SSBs or BER intermediates. To further ensure that the measured SSBs are not derived from cells that would have already enter the S phase (see the PI histograms in **Figure 6A**), we performed EdU alkaline and neutral comet assays using the same experimental design and measured tail moment in EdU-negative comets (**Figure 6D**). Again, we observed an increase in tail moment when cells were exposed to PARPi and when comet assay were performed in alkaline conditions (**Figure 6E**). No significant increase was observed when cells were treated with H2O2, suggesting that BER intermediates induced by this compound during G1 may be efficiently repaired when PARP1/2 are functional. When comet assay was performed in neutral conditions, no increase in tail moment was observed when cells were exposed to veliparib (**Figure 6F**), confirming that this inhibitor induces an accumulation of SSBs only in the DNA template at the end of G1. However, a significant increase in tail moment was observed when cells were exposed to olaparib, indicating that this inhibitor induces DSBs, and maybe also SSB, in the DNA during G1. Of note, our experimental design allows us to measure endogenous unrepaired DNA breaks generated when PARylation is inhibited only during G1, but it is also possible that these DNA breaks are generated during S phase.

**Figure 6:**
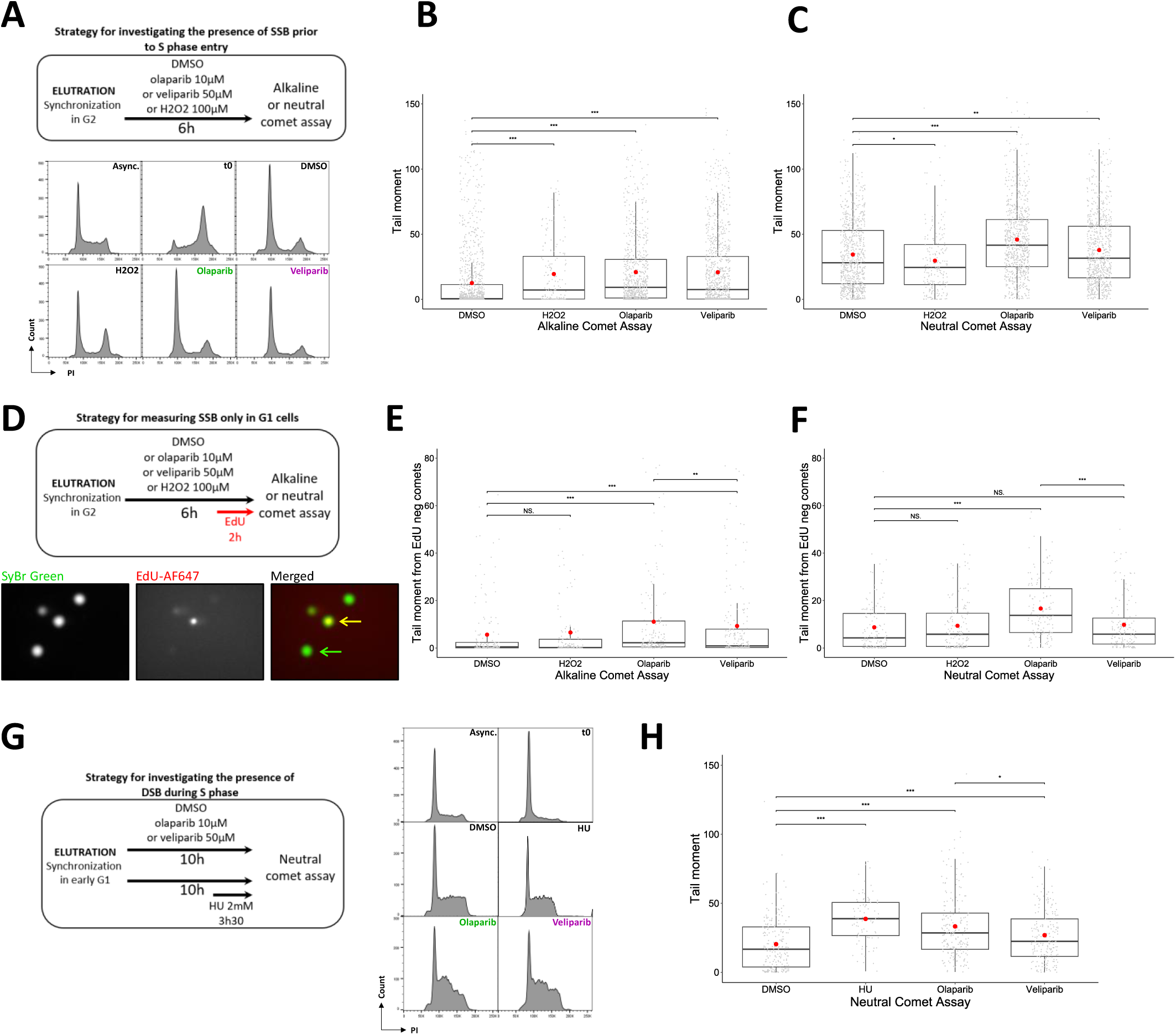
Absence of PARylation induces SSB accumulation during G1, prior to S phase entry, and DSB during S phase. **(A)** Schematic of the comet assay to investigate the presence of SSBs in DNA prior to S phase entry following inhibition of PARylation during G1. K562 cells were synchronized in G2 and treated with olaparib 10µM or veliparib 50µM for 6 hours until the end of the G1 phase. Alkaline and neutral comet assays were performed at the end of the treatment. Treatment with H2O2 100µM under the same conditions was also performed as a positive control for oxidative stress-induced DNA damage. Cell cycle profiles of samples after synchronization are displayed. **(B)** Box plots indicating the tail moment measured in comets in each condition from the alkaline comet assay. Scored comets from two independent experiments: DMSO = 1158; H2O2 = 175; olaparib = 712; veliparib = 953. Statistics: non-parametric Mann-Whitney test. ns=non-significant, * p<0.05, ** p < 0.01, *** p < 0.001. **(C)** Box plots showing the tail moment measured in comets in each condition from the neutral comet assay. Scored comets from two independent experiments: DMSO = 768; H2O2 = 211; olaparib = 725; veliparib = 734. Statistics: non-parametric Mann-Whitney test. ns=non-significant, * p<0.05, ** p < 0.01, *** p < 0.001. **(D)** Schematic of the comet assay to ensure the measurement of SSBs generated only during G1. K562 cells were synchronized in G2 and treated with olaparib 10µM, veliparib 50µM or H2O2 100µM for 6 hours until the end of the G1 phase. EdU 10µM was added to the culture medium for the last 120 minutes in order to label nascent DNA in cells that have already enter the S phase. Alkaline and neutral comet assays combined to EdU immunodetection by Click-iT chemistry were performed at the end of the treatment. An example of EdU negative (in green) and EdU positive comets (in yellow) is displayed. **(E)** Box plots indicating the tail moment measured in EdU negative comets in each condition from the alkaline comet assay. Scored comets from two independent experiments: DMSO = 179; H2O2 = 152; olaparib = 174; veliparib = 205. Statistics: non-parametric Mann-Whitney test. ns=non-significant, * p<0.05, ** p < 0.01, *** p < 0.001. **(F)** Box plots indicating the tail moment measured in EdU negative comets in each condition from the neutral comet assay. Scored comets from two independent experiments: DMSO = 140; H2O2 = 151; olaparib = 140; veliparib = 147. Statistics: non-parametric Mann-Whitney test. ns=non-significant, * p<0.05, ** p < 0.01, *** p < 0.001. **(G)** Schematic of the comet assay to investigate the presence of DSBs in DNA during S phase when PARylation is inhibited from G1 to mid S phase. K562 cells were synchronized in early G1 and treated with olaparib 10µM or veliparib 50µM for 10 hours until the mid S phase. A cell treatment with HU 2mM for the last 210 minutes (3h30) was also performed as a positive control for double-strand breaks formation. Neutral comet assay is performed at the end of the treatment. Cell cycle profiles of samples after synchronization are displayed. **(H)** Box plots indicating the tail moment measured in comets in each condition from the neutral comet assay. Scored comets from two independent experiments: DMSO = 198; HU = 85; olaparib = 204; veliparib = 196. Statistics: non-parametric Mann-Whitney test. ns=non-significant, * p<0.05, ** p < 0.01, *** p < 0.001.

To test whether these endogenous unrepaired SSBs, which are accumulated prior to S phase entry, could be transformed into one-ended DSBs during the subsequent DNA replication upon absence of PARylation, we conducted neutral comet assay on cells that had been synchronized in early G1 and cultivated with PARPi for 10 hours, until the mid S phase (**Figure 6G**). As anticipated, we observed an increase in tail moment in the neutral comet assay, indicating the generation of DSBs during S phase (**Figure 6H**). Veliparib induced fewer DSBs during S phase than olaparib (**Figure 6H**), which is concomitant with a lower induction of SSBs during G1 (**Figure 6E**) and a lower number of FANCD2 foci during S phase (**Figure 5C**). Altogether, these results confirm that an absence of PARylation induces accumulation of SSBs in the template DNA strands leading to formation of DSBs during S phase. This is consistent with the increase in γH2AX and FANCD2 foci during S phase (**Figures 2D, 2E, 5C, 5D**) indicating the formation of DNA repair foci.

### The SSBs induced by PARP inhibitors are not associated with PARP trapping

We next wanted to test whether the DNA replication defects associated with PARP inhibitors are caused by PARP1-2 trapping. Indeed, PARP1-2 trapping consists in the retention of PARP1-2 proteins on DNA, a process that is enhanced by some PARP inhibitors [74,75]. PARP inhibitors can be pro- or anti-trapping, depending on the type of inhibitor, due to their distinct effects on the allosteric conformation of the PARP1-2 proteins [76]. Olaparib has a weak trapping effect, while Veliparib does not [76]. PARP1-2 proteins have a high affinity for DNA break structures [77,78,for a review 79] and it is commonly assumed that PARP1-2 trapping on SSBs can create toxic obstacles to ongoing replicative forks and induce genetic instability, a phenomenon that is attributed to PARP inhibitor toxicity [74,75,80,for a review 81]. To detect potential PARP1-2 trapping on DNA break structures generated before or during S phase when PARylation is inhibited (**Figures 6E, 6F, 6H**), we measured the amount of the chromatin-bound PARP1 protein by chromatin fractionation and western blot. Cells were synchronized in early G1 and cultivated for 10 hours, until the mid S phase with olaparib or veliparib, in combination with or without methyl methanesulfonate (MMS), which served as a control condition for observing PARP1 trapping [74,75] (**Figure S6A**). The abundance of PARP1 in chromatin protein extracts was not increased following exposure to olaparib or veliparib, even when combined with MMS (**Figure S6B, bottom, short exposure, lanes 7 to 12**). Nevertheless, with a longer exposition time, higher molecular bands appeared in the PARP1 blot of chromatin fraction solely when cells were treated with olaparib or veliparib in combination with MMS (**Figure S6B, bottom, long exposure, lanes 11 and 12**). We assumed these bands correspond to SUMOylated and Ubiquitylated trapped PARP1 as previously described [82]. This study demonstrated that following trapping, PARP1 is SUMOylated by PIAS4 and subsequently ubiquitylated by the SUMO-targeted E3 ubiquitin ligase RNF4, which promotes the recruitment of p97 and the removal of trapped PARP1 from chromatin [82]. Therefore, we conclude that this assay was sufficiently efficient to test PARP1 trapping and that there was no evidence of PARP1 trapping detection when K562 cells were exposed to olaparib or veliparib from the G1 phase to the mid S phase (**Figure S6B, bottom, long exposure, lanes 7 to 9**). We also tested whether there was some PARP1 trapping in the conditions of the iPOND experiment, *ie.* in asynchronous cells treated with olaparib or veliparib for six hours. Once again, no evidence of PARP1 trapping was detected (**Figure S6C, right, long exposure, lanes 1 to 3**). We therefore concluded that the DNA replication defects observed when cells were exposed to PARP inhibitors for a single S phase were not primarily due to PARP1-2 trapping, although some undetectable trapping could occur, but rather to the presence of DNA breaks in the DNA template. In addition, these results suggest that the differential effects of olaparib and veliparib on replication fork progression and origin firing are not attributable to trapping.

### PARP inhibition disturbs the DNA replication timing program of some disease-associated genomic regions

We have shown that absence of PARylation induces SSBs in the replicative template DNA which are converted into one-ended DSBs during S phase (**Figure 6**). This is associated with S phase delay, ATR activation and formation of DNA repair foci (**Figures 1, 2 and 5**). Repair of one-ended DSBs at collapsed forks can involve error-prone repair pathways, especially in the absence of a replication fork coming from the opposite direction [for a review, 83]. It is therefore possible that the absence of PARylation induces genomic instability by inducing mutagenic rearrangements.

In eukaryotic genomes, the replication timing (RT) program controls the timing of activation of thousands of replication origins during the S phase. Consequently, different parts of the genome are replicated either in the early, mid or late stages of the S phase. We have previously developed the START-R suite, which allows us to analyze and compare the RT program genome-wide between two conditions, and identify genomic regions where the RT is significantly altered [58,84].

To explore in an undirect way the consequences of the potential genomic instability induced by an absence of PARylation, we performed an analysis of the RT program genome-wide of cells exposed to PARP inhibitors. Our expectation is that the genomic regions where the RT is delayed by PARPi might be regions where PARPi have induced a one-ended DSB, and might be potentially prone to mutagenic rearrangements. An alternative hypothesis is that PARPi could have reduced origin firing in these regions (**Figures 3C and 3E-G**), resulting in a delay in RT. A reduction in origin firing could lead to under-replicated regions before the end of S phase, threatening genome stability [for a review, 85]. In these two situations, genomic characterization of the specific regions where RT is delayed by PARPi could yield valuable insights into the consequences of PARP inhibition on genome stability.

We thus performed an analysis of the RT program of exponentially growing asynchronous K562 cells treated with olaparib or veliparib for 8 hours (**Figure 7A**). Briefly, cells were labelled with a pulse of BrdU before the end of the treatment with inhibitors and sorted into early and late S phase fractions. For each condition, the newly synthesized DNA of these two fractions was differently labelled and was then hybridized on microarrays. These microarrays were focused on disease-associated regions throughout the genome linked to developmental delay, intellectual disability, neuropsychiatric disorders, congenital anomalies or dysmorphic features. Thanks to the START-R suite developed in our laboratory [58], we found that olaparib and veliparib induced RT modifications of 0.81% and 3.73% of the entire genome, respectively (**Figures 7B**). We generated differential RT profiles between control and treated cells (**Figure 7C**). These percentages of genome-wide RT alterations are not as high as when factors regulating replication origins are downregulated, as we recently published [86,87]. This further suggests that PARP inhibitors do not directly affect the replication origin activity, but rather directly affect the DNA elongation process, most likely by inducing one-ended DSB formation (**Figure 6**).

**Figure 7:**
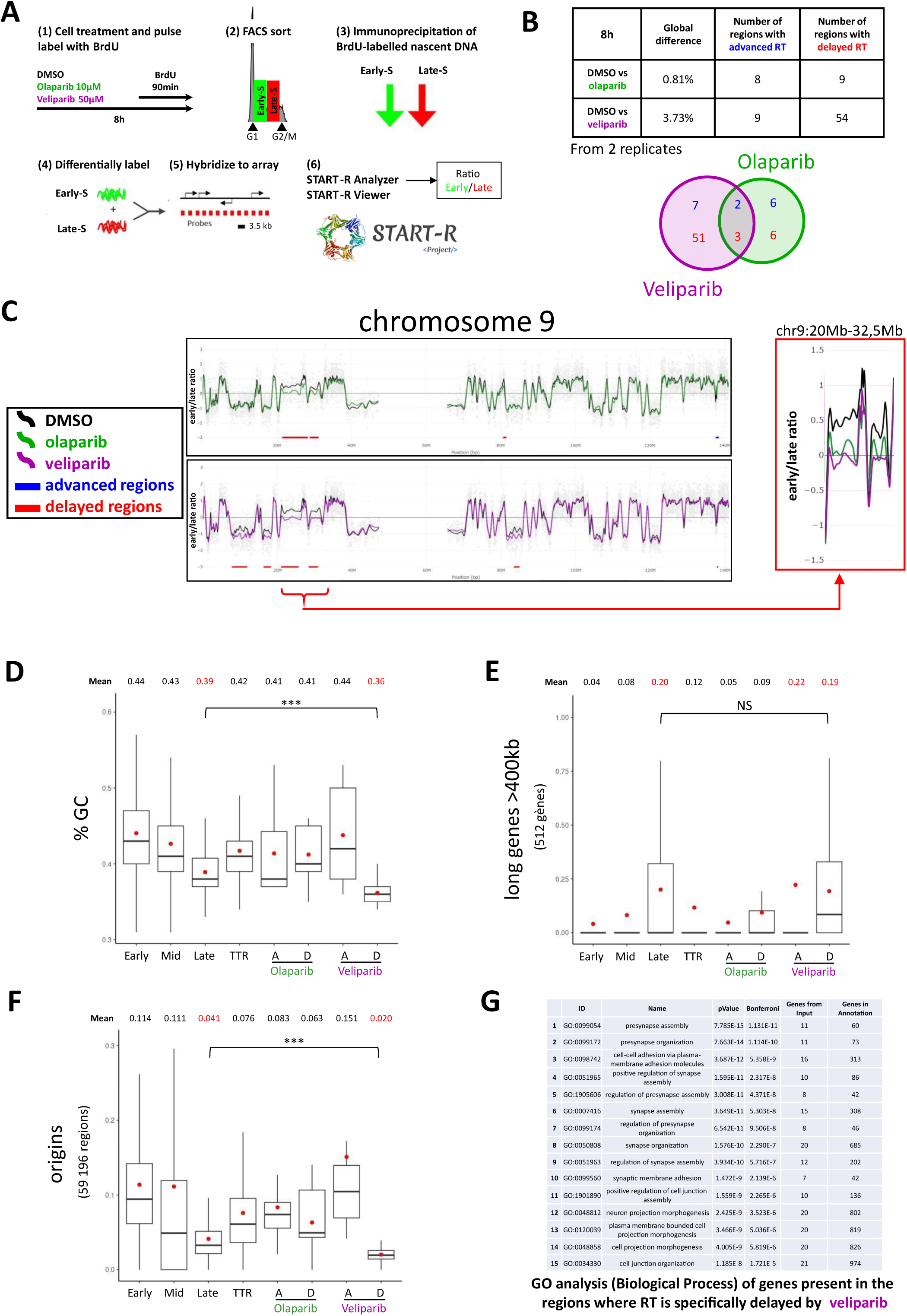
Absence of PARylation induces modification of DNA replication timing of some disease-associated genomic regions. **(A)** Schematic of the replication timing experiment. **(B)** Percentages of global difference in genome-wide replication timing between the DMSO control and the drug condition (two independent experiments were performed). The number of genomic regions where RT is either delayed or advanced by olaparib or veliparib is indicated. A Venn-diagram indicates the common regions where RT was modified by both PARPi. One of these common regions located on the chromosome 9 is highlighted in **(C)**. **(C)** Example of replication timing profiles displayed by the START-R viewer. Replicating timing profiles of untreated (DMSO, black) or treated K562 cells during 8 hours with olaparib 10µM (green) or veliparib 50µM (purple). Each grey point corresponds to a probe covering a region of the human chromosome 9 (hg19 genome assembly). The x-axis represents the chromosomal coordinates in Mb on the chromosome 9. The y-axis represents the early replicated regions (positive values) and the late replicated regions (negative values). **(D-F)** Box plots showing the percentage of GC **(D)** or the coverage of long genes > 400kb **(E)** and replication origins **(F)** in regions advanced or delayed by olaparib or veliparib. The percentage or the coverage of these genomic features in early, mid, late and TTR (Timing Transition Region) replicated regions of the K562 cell line is indicated on the left of each box plot. Mean values are indicated above the boxplots and depicted by a red dot on the boxplots. Statistics: non-parametric Mann-Whitney test. ns=non-significant, * p<0.05, ** p < 0.01, *** p < 0.001. **(G)** Gene Ontology analysis (ToppGene suite) performed on the genes located in the regions where RT is specifically delayed by veliparib.

We identified multiple genomic regions distributed across most of the chromosomes, where replication is significantly either delayed or advanced by PARP inhibitors in comparison to control cells (**Figures 7B and Supplemental Table 3**). For example, within the chromosome 9, olaparib and veliparib advanced the RT of one region (in blue) and olaparib delayed the RT of 3 regions (in red), whereas veliparib delayed the RT of 5 regions (**Figure 7C**). The mean size of regions with a RT delayed by PARP inhibitors was around 2Mb, which is larger than the ones with an advanced RT (**Figure S7A**). We found several commons regions where RT was modified both by olaparib and veliparib (**Figure 7B**). In particular, the RT of two proximal regions on the chromosome 9, that are normally replicated in early S phase, was strongly delayed by these inhibitors (**Figure 7C, zoom in the red square**). Interestingly, one of these regions corresponds to the 9p21.3 locus, which contains the *CDKN2A-B* tumor suppressor genes and the type 1 interferon (IFN) gene cluster (**Figure S7C**). Deletion of these genes is a common event that drives tumor progression in various cancers and is associated with poor prognosis [88–90]. This result suggests that absence of PARylation may lead to a genomic rearrangement in this region, potentially leading to the loss of *CDKN2A-B* and *IFN* genes and promoting tumorigenesis. Why this specific region on the chromosome 9 is more prone to spontaneous SSBs remains an open question.

Interestingly, we additionally found that veliparib had a stronger effect on RT than olaparib, with a total of 54 regions where the RT was delayed by this inhibitor (**Figure 7B**). We investigated whether these delayed regions were initially early, mid or late replicated during the S-phase. We found that veliparib delayed the RT of regions mostly replicated in late S phase in control cells, whereas olaparib delayed the RT of regions mostly replicated in early S phase (**Figure S7B**). We next characterized in detail the genomic content of the identified regions where RT was delayed by veliparib. First, the genomic sequences of these regions were extracted to measure the GC (Guanine and Cytosine) content. It is notable that the average percentage of GC in the human genome is approximately 41%. The delayed regions in the veliparib condition exhibit a markedly low GC content, falling below the level of GC content observed in the late-replicated regions of the K562 cell line (**Figure 7D**). This result suggests that PARylation inhibition by veliparib may disrupt DNA replication in AT-rich sequences, which are prone to form secondary structures and associated with heterochromatin. We then observed that these regions were characterized by an enrichment in very long genes (**Figure 7E**) and a paucity of origins (**Figure 7F**), similar to the late-replicated regions. Consistent with the limited number of origins in these regions, a paucity of CpG islands and G4 was also observed (**Figures S7D and S7E**). Most of the late-replicated regions in the human genome are difficult to replicate regions and more prone to replication stress, especially due to their paucity in replication origins [for a review 4]. Interestingly, the delayed regions in the veliparib condition exhibit some characteristics reminiscent of those observed in genomic regions known to contain Common Fragile Sites (CFSs): these late-replicated regions contain very large genes and exhibit a reduced number of origins [for a review, 91]. We therefore analyzed the presence of CFSs in these regions and found that 20 out of 54 regions where RT was delayed by veliparib were covered by CFS (**Supplemental Table 4**). Finally, to test whether the potential genomic instability associated with veliparib might affect a specific biological pathway, we performed a gene ontology analysis of the 199 genes present in the regions where RT is specifically delayed by this inhibitor (**Supplemental Table 5**). Interestingly, we found that the top biological processes were associated with the organization of synapses between neurons, with at least 34 distinct genes related to these categories (**Figure 7G and Supplemental Table 5**). Most of these genes are located on different chromosomes, without any obvious clustering, and replicated in late S phase. Some of them are very long genes (>350kb) and are covered by a CFS (**Supplemental Table 5**). Altogether, these results suggest that an absence of PARylation induced by veliparib could result in genomic instability in these genes and may lead to synaptic dysfunction, which is a major determinant of several neurodevelopmental and neurodegenerative diseases [for a review, 92].

## DISCUSSION

### Inhibition of PARylation directly disrupts DNA replication dynamics

In this study, we investigate the impact of two PARP1-2 inhibitors within a single S phase in order to analyze the primary effects of an absence of PARylation on replisome dynamics in an unperturbed S phase context. We show that both olaparib and veliparib delay the progression of S phase (**Figures 1A, B**), reduce DNA synthesis (**Figures 1D, 1E, S1C and S1D)**, induce replication stress (**Figure 2**) and genome-wide replication timing modifications of disease-associated regions (**Figure 7**) during a single S phase. These results demonstrate that a lack of PARylation has a direct impact on the replisome activity and/or progression (**Figure 8**), in addition to the mechanism of *trans*-cell cycle effects previously described [33].

**Figure 8:**
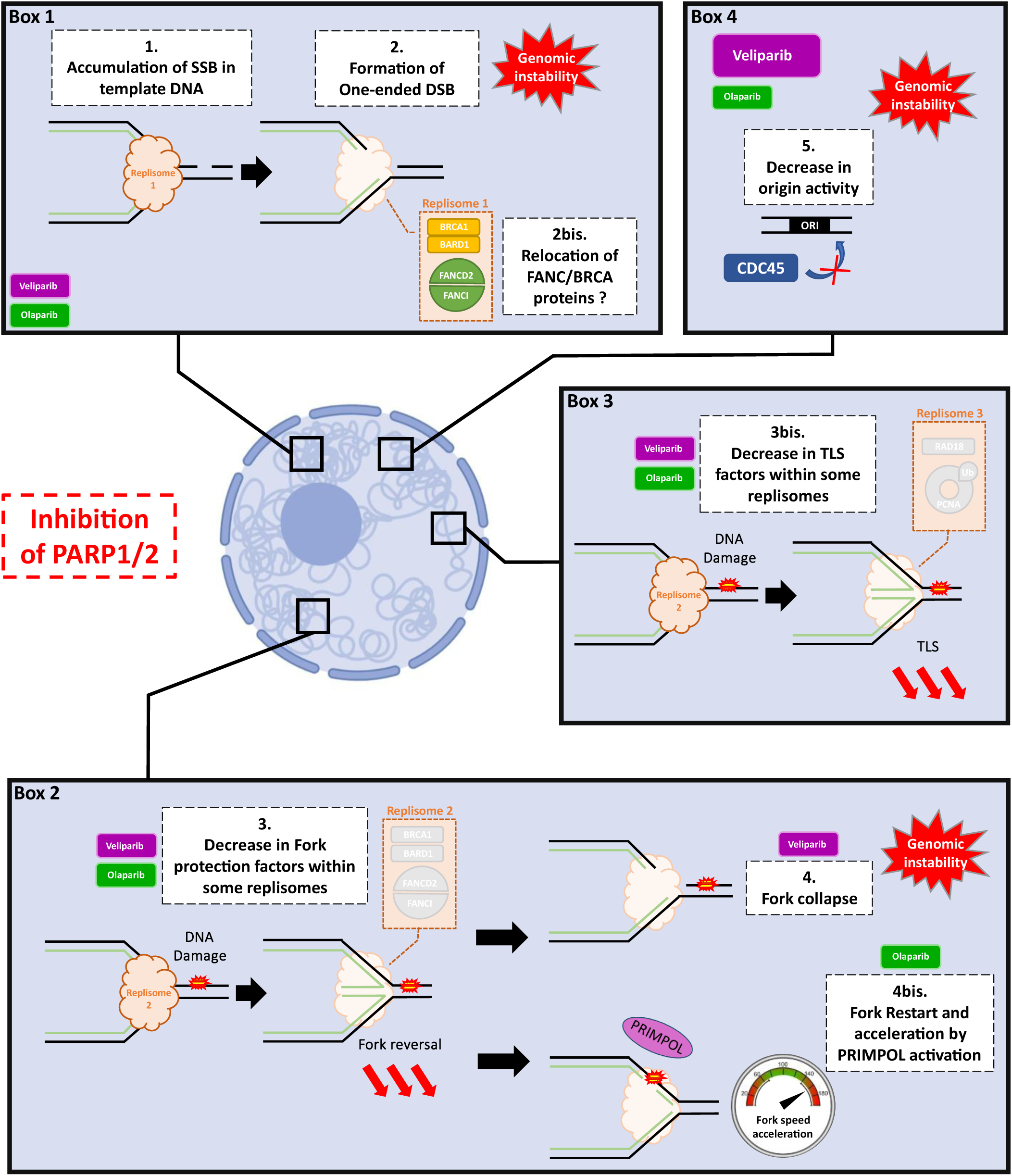
Model for the primary toxicity effects induced by PARP inhibitors during a single S phase. First, the absence of PARylation leads to the accumulation of unrepaired endogenous SSBs in the template DNA due to the inefficiency of the SSBR **(1)**. These SSBs are highly toxic lesions for the progressing replication forks during S phase, leading to the formation of one-ended DSBs and HR repair foci **(2)**. Repair of PARPi-associated one-ended DSBs may lead to mutagenic genomic rearrangements that threaten the genome integrity. Second, inhibition of PARylation during S phase remodels the protein composition of the replisome, in particular decreasing the levels of homologous recombination factors such as FANC and BRCA proteins **(3)**. This could be due to either a decrease in the recruitment of these proteins to the replisome or a relocation of these proteins from the replisome to one-ended DSB foci **(2bis)**. The decrease in FANC/BRCA proteins within the replisome may reduce fork protection and compromise fork reversal in many replicative contexts. In this situation, veliparib appears to induce persistent fork stalling, which could also lead to one-ended DSB formation due to nuclease activity **(4)**. These nuclease specific DSBs may remain unrepaired as the number of FANC/BRCA proteins is reduced in this situation, favoring genomic instability in some disease-associated regions. In contrast, olaparib appears to promote premature fork restart through PRIMPOL activity, inducing post-replicative gaps and an increase in DNA fiber length **(4bis)**. In addition, both inhibitors decrease the level of TLS factors within the replisome, which may affect the activation of this pathway **(3bis)**. Finally, by an unknown mechanism, veliparib reduces origin firing. This delays replication in many regions prone to breakage, which may result in genome instability in these regions **(5)**. Taken together, these primary effects of PARPi induce ATR activation and delay S phase progression. Some elements in the diagram were created with BioRender.com.

### Accumulation of DNA breaks ahead of replication forks is the first primary effect of PARP inhibition on DNA replication dynamics

We reveal that the absence of PARylation results in the accumulation of unrepaired SSBs, probably BER intermediates, within the DNA template (**Figures 6A-F**). These SSBs are highly toxic lesions during S phase since they are converted into one-ended DSBs by the ongoing replication forks (**Figure 6H**). This could result in the formation of HR foci during S phase to repair these one-ended DSBs (**Figures 2D-E and 5**). This accumulation of SSBs in the DNA template is expected to be the source of the replication stress observed under PARP1-2 inhibition within a single S phase (**Figures 1, 2 and 8 Box 1**). The analysis of chromatin-bound PARP1 in cells that were exposed to PARPi for a single S phase reveal an absence of trapping (**Figure S6**), indicating that the DNA replication defects observed in the presence of PARPi are likely due to unrepaired SSBs *per se*, rather than PARP1-2 trapping. The pool of damaged bases is especially pronounced in the context of elevated oxidative stress, as observed in cancerous cells [for reviews, 93,94], which may account for the relatively high level of endogenous damaged bases in the K562 cell line used in this study. The study of the genome-wide replication timing program of cells exposed to PARPi allows the investigation of the potential consequences of PARP1-2 inhibition on genome stability (**Figure 7**). We find an interesting region on the chromosome 9 where the RT is delayed by both olaparib and veliparib and that contains genes that have been implicated in tumorigenesis due to their genetic loss (**Figures 7 and S7C**). This suggests that PARPi-associated one-ended DSBs could result in genomic rearrangements in this region and promote tumorigenesis (**Figure 8 Box 1**).

### Remodeling of the replisome proteome is the second primary effect of PARP inhibition on DNA replication dynamics

Subsequently, we demonstrate that PARP inhibitors deregulate the protein composition of the replisome (**Figure 4**). Especially, absence of PARylation decreases the abundance of FANCD2-FANCI-BRCA1-BARD1 within replisome (**Figure 4C and 8 Box 2**). These results suggest that PARylation may directly or indirectly influence the presence of these proteins at replisome. It is possible that PARP activity is directly involved in the recruitment of FANC-BRCA proteins at the replisome in human cells. In accordance with this, previous studies have demonstrated that BARD1 directly interacts with PARylation through a BRCT domain, thereby enabling the BRCA1-BARD1 heterodimer to be recruited to IR-induced DSB in human cells [95], or to replisomes in mice cells [96]. Regarding FANCD2-FANCI, it has been demonstrated that the PARylation of RUNX3 enhances its interaction with the helicase BLM, which facilitates the recruitment of FANCD2 to Mitomycin C (MMC)-induced ICL sites [97]. Another study has demonstrated that olaparib partially diminishes the recruitment of FANCD2 to psoralen-induced ICL sites [98]. Interestingly, a similar result was observed in a knockdown of *PARP3* but not of *PARP1* [98]. However, the exact molecular mechanism by which FANCD2-FANCI are recruited to DNA lesions or within the replisome (**Figure 4** of this study) through PARP activity remains unknown. A PROSITE analysis revealed that FANCD2 and FANCI lack a known PARylation interacting motif (PBM, PBZ, WWE and BRCT domains were tested, data not shown). Nevertheless, the ADPriboDB 2.0 database [99,100] indicates that FANCD2 and FANCI are PARylated, which may serve to stabilize them at genomic sites where it is required, as it has already been demonstrated for other PARylated proteins [97,101,102]. A previous study demonstrated that FANCD2 is recruited within the replisome through direct interactions with MCM2-7 upon ATR signaling [53]. It is also possible that PARylation dysregulation alters this interaction. Further biochemical studies will be required to elucidate the precise mechanism by which the FANCD2-FANCI complex is recruited or stabilized within the replisome through PARP activity. Alternatively, the absence of PARylation may indirectly decrease the abundance of FANC-BRCA proteins within replisomes. Indeed, our study reveals that PARP inhibitors induce formation of FANCD2 foci during a single S phase (**Figure 5**) without impacting the total amount of FANCD2 protein (**Figures S4F-G**). These results suggest that the absence of PARylation could favor the location of FANC-BRCA proteins to HR foci rather than replisomes, or induce the relocation of these proteins from replisomes to HR foci, in order to repair PARP inhibitor-induced one-ended DSBs (**Figure 8 Boxes 1 and 2**). In both hypotheses, a decrease of FANC-BRCA proteins within replisome may impact the replisome progression by decreasing fork protection (see next section).

In addition, our iPOND experiment revealed a decrease in the abundance of RAD18 and PCNA monoubiquitination within the replisome in the presence of PARP inhibitors. RAD18 plays a pivotal role in the Translesion Synthesis (TLS) pathway, which is activated at the replisome to bypass DNA lesions and behind replication forks, to fill the post-replicative gaps generated by PRIMPOL [for recent reviews, 103,104]. Although further investigations will be required to test this hypothesis, we propose that inhibition of PARP1/2 activity may also reduce the activation of TLS at the replisome and its progression (**Figure 8 Box 3**). This could be as well a mechanism by which post-replicative gaps persist in DNA strands in the presence of PARP inhibitors [33,40].

### Absence of PARylation may decrease fork protection

Besides HR, FANCD2-FANCI and BRCA1 are implicated in replication fork protection of reversed forks, by preventing the nuclease-mediated degradation of nascent DNA at stalled forks [12,105–111]. Fork reversal is a common mechanism that is established in response to a multitude of genotoxic agents [112], which can also be observed at fork that are not directly disrupted by the presence of DNA damage [113]. Therefore, fork reversal is a mechanism that globally slows replication across the genome, allowing time for repair mechanisms to take over DNA damage. Interestingly, olaparib decreases the incidence of fork reversal, by suppressing the inhibitory effect of PARP1/2 activity on RECQ1, thereby causing a premature restart of reversed forks [11]. Considering our results, we thus propose that PARP inhibitors may exert an additional adverse effect on fork reversal by reducing the abundance of fork protective factors within the replisome, such as FANCD2-FANCI and BRCA1 (**Figure 4C**). In addition, we observe a slight increase in chromatin-bound PCNA under PARP1/2 inhibition (**Figures S4B, S4E, S5A**), an enrichment of ATAD5 and CHAF1B, a subunit of the CAF1 complex, in the olaparib replisome (**Figure 4C**). We evidence as well a decrease in the monoubiquitination of PCNA at chromatin (**Figures S5A, S5B**), probably due to a reduction in the abundance of RAD18 within the replisome in the presence of PARP inhibitors (**Figure 4C**). Interestingly, a recent study revealed that a lack of PCNA monoubiquitination leads to the retention of the protein on chromatin by preventing unloading by ATAD5, resulting in an increase in CAF1 abundance within the replisome [114]. This would ultimately result in a replicative fork with an aberrant chromatin environment, which would provide an ideal substrate for the action of nucleases during fork arrest and reversion [114]. We therefore propose that lack of PARylation during S phase may inactivate at least three (*via* RECQ1, FANC-BRCA or PCNA-ATAD5-CAF1 axes), if not all forms of replication fork protection in human cells.

### Differential effect of veliparib and olaparib on replication fork progression and origin firing

In addition, we showed that the olaparib inhibitor increases fork velocity within a single S phase (**Figures 3B**), as previously described [33,37,38]. However, the molecular mechanisms underlying the regulation of fork speed exerted by PARylation remain poorly understood. The work of Jiri Bartek’s group indicates that fork speed in human cells is negatively regulated by the interconnected PARylation and p53-p21 axis, which they have termed the Fork Speed Regulatory Network (FSRN) [37, for a review 115]. It is proposed that the acceleration of fork speed under PARP1/2 inhibition may be due in part to the deregulation of the p53-p21 expression and activity. However, as the K562 cell line is p53 negative, this p53-p21 axis is likely inactivated in these cells. Therefore, the acceleration of fork speed observed with olaparib in this cell line (**Figure 3B**) may result from a different mechanism. We show that the absence of PARylation deprives the replisome of fork protection factors, which may reduce fork reversal (**Figure 4C** and precedent section). This could provide a rationale for the activation of PRIMPOL, which is known to be active when fork reversal is suppressed [40, for a review 103] and in presence of olaparib [33,40]. Overall, this may define a molecular mechanism of faster fork restart by which the olaparib inhibitor accelerates replication forks (**Figure 8 Box 2**). In agreement with this hypothesis, a recent study have shown that PRIMPOL knockdown prevents fork acceleration induced by olaparib in human cells [39]. The fact that olaparib induces breaks in the template DNA (**Figure 6**) while accelerating replication fork speed without affecting fork symmetry (**Figure 3**) [33,37] may be counterintuitive. We would like to emphasize that the replicative machinery may certainly not be a homogeneous complex throughout the S phase and probably depending on the genomic replicative context. Therefore, it is possible that PARylation dysregulation affects differently the pool of replisomes depending on the S phase moment, the genomic sequence or the type of DNA lesions (**Figure 8 Boxes 1 and 2**). Furthermore, the DNA fiber assay, which uses short pulses of thymidine analogs, may not be the most accurate technique for detecting SSB-associated replication fork collapse events induced by PARP inhibitors, as it will only detect replication events that are actually associated with DNA synthesis. Consequently, we propose that fork asymmetry and fork acceleration may emerge concurrently in response to olaparib.

Surprisingly, the veliparib do not alter fork speed when cells are exposed to this inhibitor during a single S phase (**Figure 3B**), although it accelerates fork velocity over a longer treatment period (24 hours) [33,37,38]. Therefore, it appears that the first response of the replisome to veliparib is not to activate pathways that increase fork velocity. Moreover, to our best knowledge, there is no evidence that veliparib reduces fork reversal and induces PRIMPOL activity as olaparib. We have shown that veliparib, as olaparib, deprived replisome of fork protection factors (**Figure 4C**). In addition, veliparib induced fork asymmetry (**Figure 3C**). This suggests that the first effect of veliparib on replisome dynamics is to induce persistent fork stalling, which could lead to fork collapse in the absence of fork protective factors (**Figure 8 Box 2**). In addition, we find a unique effect of veliparib on origin activity as indicated by the IOD increase (**Figure 3D and 8 Box 4**). More strikingly, we find that veliparib delayed the RT of 54 regions across the genome (**Figure 7B**). The majority of these regions are late replicated and similar to CFS, therefore prone to breakage (**Figures 7D-F and S7D-E and Supplemental Table 4**). The fact that veliparib delays RT specifically in these regions may be due to its ability to reduce origin firing in regions already poor in origins. It may also be caused by the persistence of one-ended DSB and/or fork stalling, which cannot be efficiently repaired by a replication fork coming from the opposite direction, due to the lack of active origins in these regions. Importantly, we observe that these regions are enriched in genes associated with synapse organization (**Figure 7G and Supplemental Table 5**), which unveils that the genomic instability caused by veliparib could result in synaptopathies and neurological disorders. These results are consistent with the existence of human neurological diseases associated with mutations in SSBR components [for a review, 73], and suggest that genomic instability induced by a lack of PARP1-2 activity could disrupt the expression of some neurological genes.

A remaining key question is why olaparib and veliparib have different effects on DNA replication. A meta-analysis of the potency of olaparib (log IC_50_) *versus* the potency of veliparib in 1000 different cell lines revealed a very weak correlation between both molecules [for a review, 116]. This confirms that these two inhibitors have diverse mechanisms of action and may be indicative of differences in their ability to target PARP1-2 and potentially other proteins. According to our results (**Figure S6**), these differences do not seem to be related to the pro- or anti-trapping effects of these inhibitors. As olaparib and veliparib impact the allosteric conformation of PARP1 and 2 in various ways [76], it is also possible that these two inhibitors have different effects on the direct interactions of PARP1 and 2 with other proteins. Although we do not have any molecular explanation for the distinct effects of olaparib and veliparib, our findings provide interesting characteristics regarding the different outcomes of these two inhibitors on replication fork progression and genomic instability.

In conclusion, our study highlights the importance of maintaining PARylation during S phase to ensure DNA integrity and the accurate regulation of the replisome proteins throughout the whole process of DNA replication. We report the common and unique primary toxic effects of two PARP inhibitors, olaparib and veliparib, during S phase (**Figure 8**). Both inhibitors appear to commonly alter the balanced distribution of the FANC-BRCA pool between fork protection and DSB repair foci. However, these two inhibitors then seem to drive different pathways at replication forks and/or at replication origins. Overall, these findings could be exploited in the clinic, in order to help develop therapeutic strategies for the treatment of HR-deficient cancer cells.

## ACKNOWLEDGEMENTS

We acknowledge the ProteoSeine core facility of the Institut Jacques Monod (supported by Université Paris Cité, IBiSA, région Ile de France and CNRS) for producing the LC-MS/MS data and performing the related statistical analyses. We acknowledge the ImagoSeine core facility of the Institut Jacques Monod, member of the National Infrastructure France-BioImaging (https://ror.org/01y7vt929) supported by the French National Research Agency (ANR-24-INBS-0005 FBI BIOGEN) and GIS-IBiSA and the support of La Ligue contre le Cancer (R03/75-79), for providing cell analyzers, producing cell sorting and performing cleaning of the flow cytometry data. We thank F. Rosseli for sharing FANCD2 antibodies and C. Ribeyre for sharing biotin-TEG-azide. We also thank J. Rageul, A. Quinet and G. Guilbaud for critical comments on the manuscript.

## AUTHOR CONTRIBUTIONS

Lina-Marie Briu: Conceptualization, Data curation, Formal analysis, Investigation, Validation, Visualization, Writing – original draft, Writing – review & editing. Chrystelle Maric: Conceptualization, Investigation, Formal analysis, Validation, Writing – review & editing. Nicolas Valentin (ImagoSeine): Investigation, Formal analysis, Resources. Nicolas Panara: Investigation. Guillaume Chevreux (ProteoSeine): Investigation, Formal analysis, Resources. Giuseppe Baldacci: Conceptualization, Writing – review & editing. Jean-Charles Cadoret: Conceptualization, Investigation, Funding acquisition, Supervision, Validation, Writing – review & editing.

## SUPPLEMENTARY DATA

**Supplemental table 1**: list of the 686 proteins enriched in the K562 replisome under physiological conditions identified by iPOND-MS. Data are sorted by the “peptide LFQ” value.

**Supplemental table 2**: list of the proteins which are enriched or depleted in the K562 replisome when cells are treated with olaparib (sheet1) or veliparib (sheet 2). Data are sorted by the “Fold change” value.

**Supplemental table 3**: coordinates of the genomic regions where RT is advanced or delayed by olaparib (sheet 1) or veliparib (sheet 2).

**Supplemental table 4**: list of the 54 genomic regions where RT is delayed by veliparib. The coordinates of the CFS that overlap 20 of these regions are indicated.

**Supplemental table 5**: list of the genes located in the 51 regions where RT is specifically delayed by veliparib. The table is compound of 3 sheets: the 432 outputs from the intersect between the coordinates of the 51 regions and the coordinates of the file from the UCSC table browser RefSeq Genes database (sheet 1); the 199 gene symbols obtained after elimination of duplicates from sheet 1 (sheet 2); the 34 genes related to synapse organization obtained by GO analysis (sheet 3).

## DATA AVAILABILITY

Replication timing data are deposited in GEO Database with accession number GSE289208. The following secure token has been created to allow review of record GSE289208 while it remains in private status: ozijmmiizvqbxev. https://www.ncbi.nlm.nih.gov/geo/query/acc.cgi?acc=GSE289208

iPOND-MS/MS data are deposited in PRIDE Database with accession number PXD060870. The following secure token has been created to allow review of record PXD060870 while it remains in private status: B4isN4kLncAB.

https://www.ebi.ac.uk/pride/login

Alternatively, reviewer can access the dataset by logging in to the PRIDE website using the following account details:

Username

Password: yDSdNR5wbp6O

Concerning the flow cytometry data, we would like to inform you that the FlowRepository database is currently experiencing some difficulties. This is causing delays in the data submission process. We apologize for this inconvenience and hope that it will be resolved as quickly as possible.

